# Optimization of single dose VSV-based COVID-19 vaccination in hamsters

**DOI:** 10.1101/2021.09.03.458735

**Authors:** Kyle L. O’Donnell, Chad S. Clancy, Amanda J. Griffin, Kyle Shifflett, Tylisha Gourdine, Tina Thomas, Carrie M. Long, Wakako Furuyama, Andrea Marzi

## Abstract

The ongoing COVID-19 pandemic has resulted in global effects on human health, economic stability, and social norms. The emergence of viral variants raises concerns about the efficacy of existing vaccines and highlights the continued need the for the development of efficient, fast-acting, and cost-effective vaccines. Here, we demonstrate the immunogenicity and protective efficacy of two vesicular stomatitis virus (VSV)-based vaccines encoding the SARS-CoV-2 spike protein either alone (VSV-SARS2) or in combination with the Ebola virus glycoprotein (VSV-SARS2-EBOV). Intranasally vaccinated hamsters showed an early CD8^+^ T cell response in the lungs and a greater antigen-specific IgG response, while intramuscularly vaccinated hamsters had an early CD4^+^ T cell and NK cell response. Intranasal vaccination resulted in protection within 10 days with hamsters not showing clinical signs of pneumonia when challenged with three different SARS-CoV-2 variants. This data demonstrates that VSV-based vaccines are viable single-dose, fast-acting vaccine candidates that are protective from COVID-19.

## Introduction

Severe acute respiratory syndrome coronavirus-2 (SARS-CoV-2) has emerged as a novel, highly infectious, respiratory CoV and is the causative agent of Coronavirus disease 2019 (COVID-19), first described in the city of Wuhan in Hubei province in China (Song et al., 2020). The World Health Organization declared the SARS-CoV-2 pandemic a Public Health Emergency of International Concern on January 30^th^ 2020 (WHO, 2020). Clinically, COVID-19 can lead to respiratory distress and, in some cases, respiratory failure (Guan et al., 2020). CoVs are enveloped, single-stranded positive-sense RNA viruses with a 30 kb genome and 5 open reading frames including the four major structural proteins: spike (S), envelope, membrane, and nucleocapsid (N). The S mediates binding of SARS-CoV-1 and SARS-CoV-2 to angiotensin-converting enzyme 2 (ACE2) on the surface of various cell types including epithelial cells of the respiratory tract (Hamming et al., 2004, Letko et al., 2020, Walls et al., 2020). The COVID-19 pandemic mandated the development of a vaccine to be a global priority (Chen et al., 2020, Holshue et al., 2020, Li et al., 2020, Wu et al., 2020, Zhou et al., 2020). Due to the mutagenic nature in the replication of RNA viruses, new viral variants of concern (VOC) have emerged to dominate the pathogenic landscape. Two of the first variants that emerged were B.1.1.7 (UK; alpha variant) and B.1.351 (South Africa, SA; beta variant). B.1.1.7 acquired 23 mutations including N501Y within the S shown to increase binding affinity to the ACE2 receptor (Rambaut et al., 2020, Faria et al., 2021). B.1.351 harbors similar mutations such as the N501Y, in addition to K417N and E484K which may reduce the efficacy of existing countermeasures (Chen et al., 2021, Liu et al., 2021, Wibmer et al., 2021).

An ideal vaccine candidate would be safe, effective, rapidly deployable, require only a single immunization, and retain efficacy against multiple variants. Currently, vaccine candidates express the trimeric SARS-CoV-2 S as the primary antigen. One mRNA-based vaccine and an adenovirus-based vector have received emergency use authorization by the Food and Drug Administration (FDA) in the United States, and another mRNA vaccine recently received full FDA approval (FDA, 2021). All utilize the S as the primary antigen and elicit T cell and antigen-specific IgG responses (Corbett et al., 2020, Vogel et al., 2020, Sadoff et al., 2021). The mRNA vaccine by Pfizer received. The route of vaccination can greatly influence the local immune environment at the site of vaccination. A study comparing intramuscular (IM) and intranasal (IN) vaccination of mice with a chimpanzee adenoviral vector-based COVID-19 vaccine revealed an increase in stimulation of local mucosal immunity. Local mucosal immunity was improved after IN vaccination demonstrated by antigen specific IgA and lung resident T cell generation (Hassan et al., 2020). Benefits of IN vaccination have been demonstrated for other adenoviral vector vaccines as well as subunit vaccines, which lead to the exploration of optimal route of vaccination in this study (An et al., 2020, van Doremalen et al., 2021, King et al., 2021).

The recombinant vesicular stomatitis virus (VSV) vaccine platform has previously been used for multiple viral pathogens such as Ebola, Nipah, and Lassa (Safronetz et al., 2015, Mire et al., 2019, Marzi et al., 2011). We developed two VSV-based vaccines for SARS-CoV-2: a monovalent and a bivalent vaccine construct. The monovalent construct expresses the S of SARS-CoV-2 (VSV-SARS2) with a cytoplasmic tail deletion, which has been previously described (Dieterle et al., 2020). Recently, a similar VSV-based vaccine expressing the full-length S demonstrated protective efficacy against COVID-19 in Syrian golden hamsters challenged 23 days after IM vaccination (Yahalom-Ronen et al., 2020). The bivalent vaccine co-expresses the full-length S and the Ebola virus (EBOV) GP (VSV-SARS2-EBOV). The VSV vaccine platform displays several advantages to other similar approaches. VSV-based vaccines have been shown to produce a robust and rapid immune response to the encoded antigen(s) after a single immunization. Other viral vector vaccines have the problem of preexisting immunity; with VSV preexisting immunity would be directed primarily against the glycoprotein, which is not present in this system (Fathi et al., 2019). The time to immunity has been demonstrated to be 7 to 10 days for a number of pathogens, greatly reducing the time needed between vaccination and protection (Fathi et al., 2019, Marzi et al., 2015). Multiple routes of vaccination have been shown to be efficacious utilizing VSV-based vaccines, such as IM and IN (Brown et al., 2011, Fathi et al., 2019, Furuyama et al., 2020, Henao-Restrepo et al., 2017, Marzi et al., 2015). Previously, we determined the efficacy of IM and IN vaccination of nonhuman primates (NHP) with VSV-SARS2-EBOV. The study demonstrated that IM vaccination resulted in superior protective efficacy with a short time to challenge, however, IN vaccination might be similar with a longer time between vaccination and challenge (Furuyama et al., 2021). These unique attributes - robust immune stimulation and short time to immunity - make VSV an attractive viral vector vaccine platform for SARS-CoV-2.

Syrian golden hamsters have previously been established as a model system for SARS-CoV-2 recapitulating respiratory disease (Imai et al., 2020, Rosenke et al., 2020). When IN challenged, these animals develop moderate broncho-interstitial pneumonia with peak viral replication in the lungs 3 days post challenge (DPC) resolving by day 10. Peak histopathologic lesions in the lungs have been observed between 3-to 5-DPC (Rosenke et al., 2020). In this Syrian golden hamster study, we sought to determine the humoral and cellular immunogenicity over time in response to two VSV-based SARS-CoV-2 vaccines through both IN- and IM-vaccination routes at two challenge timepoints. We show that both vaccines offer protective immunity against multiple viral variants in the Syrian golden hamster model.

## Materials & Methods

### Ethics statement

All infectious work with SARS-CoV-2 was performed in the high-containment laboratories at the Rocky Mountain Laboratories (RML), Division of Intramural Research, National Institute of Allergy and Infectious Diseases, National Institutes of Health. RML is an institution accredited by the Association for Assessment and Accreditation of Laboratory Animal Care International (AAALAC). All procedures followed standard operating procedures (SOPs) approved by the RML Institutional Biosafety Committee (IBC). Animal work was performed in strict accordance with the recommendations described in the Guide for the Care and Use of Laboratory Animals of the National Institute of Health, the Office of Animal Welfare and the Animal Welfare Act, United States Department of Agriculture. The studies were approved by the RML Animal Care and Use Committee (ACUC). Procedures were conducted in animals anesthetized by trained personnel under the supervision of veterinary staff. All efforts were made to ameliorate animal welfare and minimize animal suffering; food and water were available ad libitum.

### Animal study

Two hundred and fifty Syrian golden hamsters (5-8 weeks of age; male and female) were used in this study. The hamsters were randomly selected into groups as shown in table S2. On the day of vaccination hamsters received a single dose of 1×10^5^ PFU of VSV-SARS2-EBOV or VSV-SARS2 by the IM (thigh) or IN route. Control animals received the same dose of a control vaccine (VSV-EBOV) by either the IM or IN route. On days 3, 10, and 38 animals were euthanized for sample collection to analyze vaccine immunogenicity. For efficacy studies with 28 and 10 days between vaccination and challenge animals received the same vaccine dose by the above mentioned routes. On day 0, all hamsters animals were challenged IN with 1× 10^5^ TCID_50_ SARS-CoV-2 as previously described (Rosenke et al., 2020). On 4 DPC, all animals were euthanized for sample collection.

### Cells and Viruses

Huh7 and VeroE6 cells were grown at 37°C and 5% CO_2_ in Dulbecco’s modified Eagle’s medium (DMEM) (Sigma-Aldrich, St. Louis, MO) containing 10% fetal bovine serum (FBS) (Wisent Inc., St. Bruno, Canada), 2 mM L-glutamine (Thermo Fisher Scientific, Waltham, MA), 50 U/mL penicillin (Thermo Fisher Scientific), and 50 μg/mL streptomycin (Thermo Fisher Scientific). BHK-T7 (baby hamster kidney) cells expressing T7 polymerase were grown at 37°C and 5% CO_2_ in minimum essential medium (MEM) (Thermo Fisher Scientific) containing 10% tryptose phosphate broth (Thermo Fisher Scientific), 5% FBS, 2 mM L-glutamine, 50 U/mL penicillin, and 50 μg/mL streptomycin. Ancestral SARS-CoV-2 isolate nCoV-WA1-2020 (MN985325.1) (Harcourt et al., 2020), SARS-CoV-2 isolate B.1.1.7 (hCOV_19/England/204820464/2020), or SARS-CoV-2 isolate B.1.351 (hCoV-19/South African/KRISP-K005325/2020) were used for the animal challenge studies and neutralization testing. The following reagent was obtained through BEI Resources, NIAID, NIH: Severe Acute Respiratory Syndrome-Related Coronavirus 2, Isolate hCoV-19/England/204820464/20200, NR-54000, contributed by Bassam Hallis. SARS-CoV-2 B. 1.351 was obtained with contributions from Dr. Tulio de Oliveira and Dr. Alex Sigal (Nelson R Mandela School of Medicine, UKZN). All viruses were grown and titered on Vero E6 cells, and sequence confirmed.

### Generation of VSV-based vaccine candidates

The SARS-CoV-2 S open reading frame was PCR-amplified from an expression plasmid encoding the codon-optimized (human) gene based on GenBank accession number MN908947 which was kindly provided by Vincent Munster (NIAID). Full-length SARS-CoV-2 S was cloned into the pATX-VSV-EBOV plasmid upstream of the EBOV-Kikwit GP resulting in VSV-SARS2-EBOV (Fig. S1A) following a previously successful strategy (Tsuda et al., 2011). The cytoplasmic tail deletion was introduced by PCR and was cloned into the pATX-VSV plasmid resulting in VSV-SARS2. The replication competent recombinant VSV was recovered in BHK-T7 cells as described previously (Emanuel et al., 2018). VSV-SARS2-EBOV was propagated in Huh7 cells. The complete sequence of the virus was confirmed by Sanger sequencing. The titer of the virus stock was quantified using standard plaque assay on VeroE6 cells.

### Growth kinetics

VeroE6 cells were grown to confluency in a 12-well plate and infected in triplicate with VSVwt, VSV-EBOV, VSV-SARS2, or VSV-SARS2-EBOV at a multiplicity of infection of 0.01. After 1 h incubation at 37°C, cells were washed three times with plain DMEM, and covered with DMEM containing 2% FBS. Supernatant samples were collected at 0, 6, 12, 24, 48, 72, and 96 hours post infection and stored at −80 °C. The titer of the supernatant samples was determined performing TCID_50_ assay on VeroE6 cells as previously described (Emanuel et al., 2018).

### Western blot analysis

Supernatant samples containing VSV were mixed 1:1 with sodium dodecyl sulfate-polyacrylamide (SDS) gel electrophoresis sample buffer containing 20% β-mercaptoethanol and heated to 99 °C for 10 min. SDS-PAGE and transfer to Trans-Blot polyvinylidene difluoride membranes (Bio-Rad Laboratories) of all samples was performed as described elsewhere (Furuyama et al., 2020). Protein detection was performed using anti-SARS-CoV-2 S RBD (1:1000; Sino Biological) or anti-EBOV GP (ZGP 12/1.1, 1 μg/ml; kindly provided by Ayato Takada, Hokkaido University, Japan) or anti-VSV M (23H12, 1:1000; Kerafast Inc.). After horse-radish peroxidase (HRP)-labeled secondary antibody staining using either anti-mouse IgG (1:10,000) or anti-rabbit IgG (1:5000) (Jackson ImmunoResearch), the blots were imaged using the SuperSignal West Pico chemiluminescent substrate (Thermo Fisher Scientific) and an iBright™ CL1500 Imaging System (Thermo Fisher Scientific).

### RNA extraction and RT-qPCR

Nasal swab samples were extracted using the QIAamp Viral RNA Mini Kit (Qiagen) according to manufacturer specifications. Tissues, a maximum of 30 mg each, were processed and extracted using the RNeasy Mini Kit (Qiagen) according to manufacturer specifications. One step RT-qPCR for genomic viral RNA was performed using specific primer-probe sets and the QuantiFast Probe RT-PCR +ROX Vial Kit (Qiagen), in the Rotor-Gene Q (Qiagen) as described previously (van Doremalen et al., 2020). Five μL of each RNA extract were run alongside dilutions of SARS-CoV-2 standards with a known concentration of RNA copies.

### Enzyme-linked immunosorbent assay

Serum samples from SARS-CoV-2 infected animals were inactivated by γ-irradiation and used in BSL2 according to IBC-approved SOPs. NUNC Maxisorp Immuno plates were coated with 50 μl of 1 μg/mL of recombinant SARS-CoV-2 S (S1+S2), SARS-CoV-2 RBD (Sino Biological) or EBOV GP at 4°C overnight and then washed three times with PBS containing 0.05% Tween 20 (PBST). The plates were blocked with 3% skim milk in PBS for 3 hours at room temperature, followed by three additional washes with PBST. The plates were incubated with 50 μl of serial dilutions of the samples in PBS containing 1% skim milk for 1 hour at room temperature. After three washes with PBST, the bound antibodies were labeled using 50 μl of 1:2,500 peroxidase anti-hamster IgG (H+L) (SeraCare Life Sciences) diluted in 1% skim milk in PBST. After incubation for 1 h at room temperature and three washes with PBST, 50 μl of KPL ABTS peroxidase substrate solution mix (SeraCare Life Sciences) was added to each well, and the mixture was incubated for 30 min at room temperature. The optical density (OD) at 405 nm was measured using a GloMax® explorer (Promega) plate reader. The OD values were normalized to the baseline samples obtained with naïve hamster serum and the cutoff value was set as the mean OD plus standard deviation of the blank.

### Flow cytometry

Hamster PBMCs were isolated from ethylene diamine tetraceticacid (EDTA) whole blood by overlay on a Histopaque®-1077 density cushion and separated according to manufacturer’s instructions. Tissues were processed into single cell suspensions as described previously (Barrigan et al., 2013). Cells were stimulated for 6 hours with media alone, cell stimulation cocktail (containing PMA-Ionomycin, Biolegend), 1μg/ml SARS-CoV-2 S peptide pool, or Lassa virus (LASV) GPC peptide pool together with 5μg/ml Brefeldin A (Biolegend). Following surface staining with Live/Dead-APC/Cy7, CD4-Alexa700, CD8-FITC, CD94-BV421 and CD69-PeCy7, B220-BV605, CD11b-PerCPCy5.5, and Ly6G-APC (all Biolegend) cells were fixed with 4% paraformaldehyde (PFA). Sample acquisition was performed on a FACSSymphony-A5 (BD), and data analyzed in FlowJo V10. Cell populations were identified by initially gating on Live/Dead negative, doublet negative (SSC-H vs SSC-A). Activation positive responses are presented after subtraction of the background responses detected in the LASV GPC peptide pool-stimulated samples.

### Virus neutralization assay

The day before this assay, VeroE6 cells were seeded in 96-well plates. Serum samples were heat-inactivated for 30 min at 56°C, and 2-fold serial dilutions were prepared in DMEM with 2% FBS. Next, 100 TCID_50_ of SARS-CoV-2 were added and the mixture was incubated for 1 hour at 37°C and 5% CO_2_. Finally, media was removed from cells and the mixture was added to VeroE6 cells and incubated at 37°C and 5% CO_2_ for 6 days. CPE was documented, and the virus neutralization titer was expressed as the reciprocal value of the highest dilution of the serum which inhibited virus replication (no CPE)(van Doremalen et al., 2020).

### Histology and immunohistochemistry

Tissues were fixed in 10% neutral buffered formalin with two changes, for a minimum of 7 days. Tissues were placed in cassettes and processed with a Sakura VIP-6 Tissue Tek, on a 12-hour automated schedule, using a graded series of ethanol, xylene, and ParaPlast Extra. Embedded tissues are sectioned at 5 um and dried overnight at 42 degrees C prior to staining. Specific anti-CoV immunoreactivity was detected using Sino Biological Inc. SARS-CoV/SARS-CoV-2 N antibody (Sino Biological cat#40143-MM05) at a 1:1000 dilution. The secondary antibody was the Vector Laboratories ImPress VR anti-mouse IgG polymer (cat# MP-7422). The tissues were then processed for immunohistochemistry using the Discovery Ultra automated stainer (Ventana Medical Systems) with a ChromoMap DAB kit (Roche Tissue Diagnostics cat#760–159). All tissue slides were evaluated by a board-certified veterinary pathologist, a representative low (20x) and high (200x) magnification photomicrograph of lung from each group was selected. Lung sections were analyzed for evidence of interstitial pneumonia and assigned the following scores: 0 normal, 1 minimal, 2 mild, 3 moderate, 4 severe.

### Statistical analyses

All statistical analysis was performed in Prism 8 (GraphPad). The serology, cellular response, RNA levels, titers and growth kinetics were examined using two-way ANOVA with Tukey’s multiple comparisons to evaluate statistical significance at all timepoints. Two-tailed Mann-Whitney or Wilcoxon tests were conducted to compare differences between groups for all other data. A Bonferroni correction was used to control for type I error rate where required. Statistically significant differences are indicated as p<0.0001 (****), p<0.001 (***), p<0.01 (**) and p<0.05 (*).

## Results

### Vaccine construction and characterization

The VSV full-length plasmid encoding the EBOV-Kikwit GP, the primary antigen for the approved EBOV vaccine, was used as the parental vector to construct the COVID-19 vaccines. First, we generated a bivalent VSV construct co-expressing the EBOV GP and SARS-CoV-2 S (VSV-SARS2-EBOV) by adding the full-length codon-optimized SARS-CoV-2 S upstream of the EBOV GP into the existing VSV vector (Fig. S1A). Second, we generated a monovalent VSV construct by replacing the EBOV GP with the SARS-CoV-2 S which contains a cytoplasmic tail deletion previously described (Case et al., 2020, Dieterle et al., 2020). Both constructs were recovered from plasmid following previously established protocols (Emanuel et al., 2018). Expression of both antigens, SARS-CoV-2 S and EBOV GP, was confirmed by Western blot analysis of the VSV particles in cell supernatant (Fig. S1B). Next, we performed viral growth kinetics and found that VSV-SARS2-EBOV replicated with similar kinetics and had comparable endpoint titers as the parental VSV-EBOV (Fig. S1C). In contrast, VSV-SARS2 showed an attenuated growth curve, and the endpoint titer was significantly lower compared to the VSV-SARS2-EBOV, potentially impacting vaccine production.

### VSV-based vaccines elicit antigen-specific humoral responses

Groups of Syrian golden hamsters (Table S1) were vaccinated with 1×10^5^ plaque forming units (PFU) either IM or IN with VSV-EBOV (control), VSV-SARS2, or VSV-SARS2-EBOV. Blood samples were collected at 3, 10, 21, and 38 days post vaccination (DPV). The humoral immune response to vaccination was examined by enzyme-linked immunosorbent assay (ELISA) using recombinant full-length S, recombinantly expressed S receptor binding domain (RBD), and recombinantly expressed EBOV GP. S-specific IgG antibodies were detected 10 DPV in the sera of both the IM- and IN-vaccinated groups for VSV-SARS2 and VSV-SARS2-EBOV (Fig. 1A, B) with antibody titers significantly higher in the VSV-SARS-EBOV IM group at 21 and 38 DPV (Fig. 1A). Hamsters in the control groups (VSV-EBOV-vaccinated) had no detectable S-specific or S RBD-specific IgG (Fig. 1A-D). Similar to the S-specific IgG response, all animals vaccinated with VSV-SARS2 and VSV-SARS2-EBOV developed measurable antibody titers to the S RBD, independent of vaccination route (Fig. 1C, D). RBD-specific antibody titers were significantly increased in the VSV-SARS2-EBOV IN-vaccinated animals at 10 DPV only (Fig. 1D). Significantly higher antibody titers for EBOV GP were not detected between VSV-EBOV and VSV-SARS2-EBOV except for 21 DPV in the IN group only (Fig. 1E, F). Antibody functionality was assessed by SARS-CoV-2 neutralization and resulted in no significant difference between the IM-vaccinated groups (Fig. 1G, H). Only VSV-SARS2-EBOV IN-vaccinated animals had a significantly higher neutralization titer compared to VSV-SARS2 at 21 DPV (Fig. 1E). Overall, VSV-SARS2-EBOV elicited a more robust and durable antigen-specific humoral response in hamsters particularly after IN administration.

**Figure 1.**
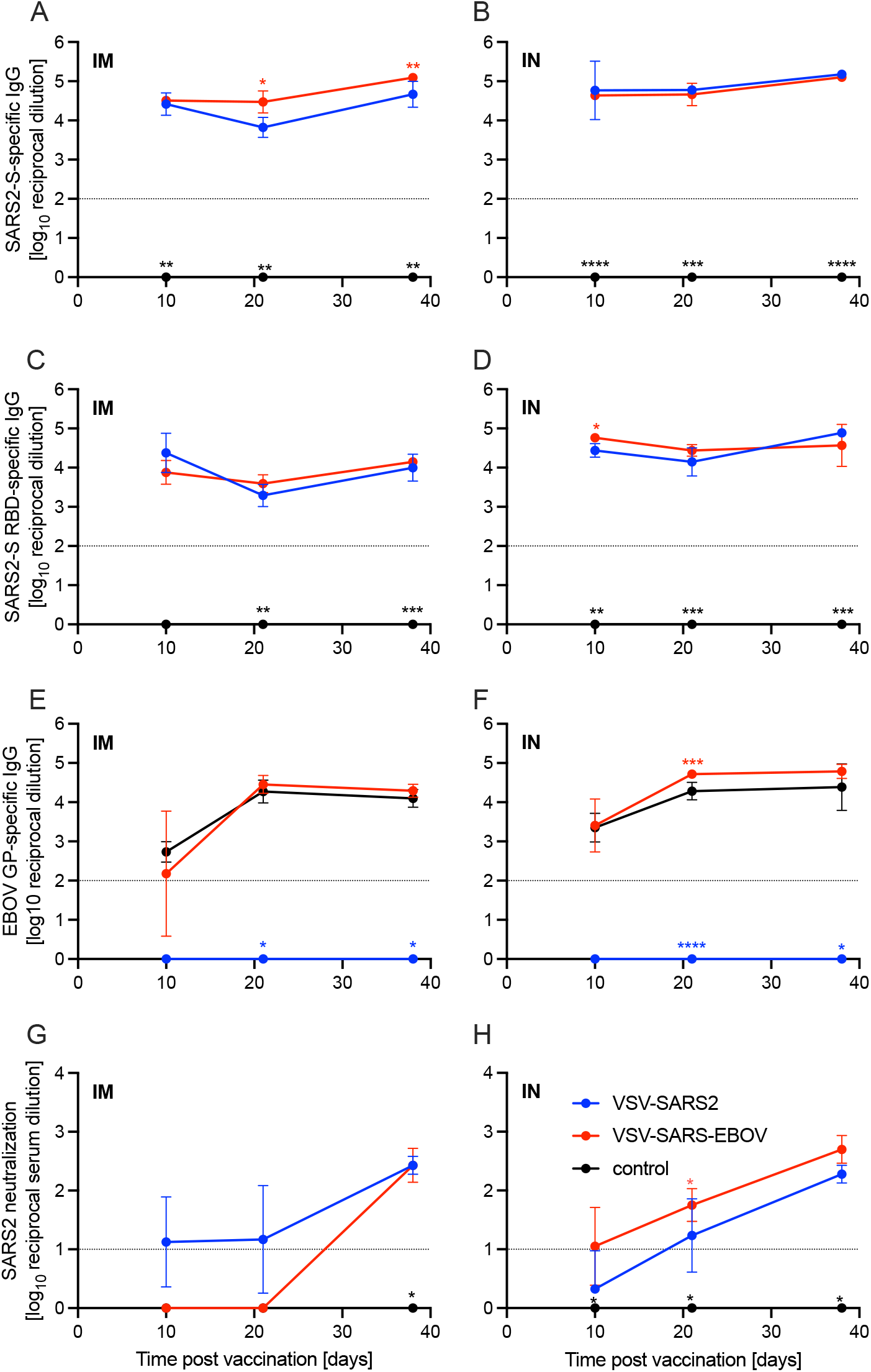
Immunogenicity humoral immune response. Serum samples were collected at multiple time points after vaccination to determine the progression of the antigen-specific antibody response by ELISA. **(A, B)** SARS-CoV-2 S-specific IgG. **(C, D)** SARS-CoV-2 S receptor binding domain (RBD)-specific IgG. **(E, F)** Ebola virus glycoprotein (EBOV GP)-specific IgG. Geometric mean and geometric SD are depicted. Statistical significance as determined by two-way ANOVA with Tukey’s multiple comparison is indicated as p<0.0001 (****), p<0.001 (***), p<0.01 (**), and p<0.05 (*).

### VSV-based vaccines induce limited cellular response

Given the potential role of cellular immunity to contribute to immune protection as seen with SARS-CoV-1 and Middle East respiratory syndrome, we sought to use flow cytometry to characterize the cellular populations involved (Channappanavar et al., 2014a, Channappanavar et al., 2014b, Zhao et al., 2010). Cellular immunology is a particular challenge in the hamster model due to the limited number of reagents available. A panel of mouse- and rat-specific flow cytometry antibodies was screened for cross-reactivity to characterize multiple cellular populations (Table S2). After we identified 7 antibodies that reacted in our initial tests, samples collected on 3, 10, and 38 DPV were used to monitor the change in cellular phenotypes over time. Single cell suspensions were created for the lungs, spleen, and peripheral blood mononuclear cells (PBMCs) and labeled for CD4, CD8, and CD69 to characterize activated T cell populations, CD94 to identify natural killer (NK) cells, B220 to stain for B cells, as well as C11b and Ly6G to identify neutrophil populations. We detected a greater percentage of activated CD4^+^ T cells in IM-vaccinated hamsters 3DPV, however, overall CD4^+^ T cell responses peaked in the VSV-SARS2-EBOV IN group 10 DPV (Fig. 2A, B). There was more overall CD8^+^ T cell stimulation on 3 and 10 DPV in the IN groups, but significantly more activated lung CD8^+^ T cells were produced in the same time frame for the IM-vaccinated animals (Fig. 2C, D). IM-vaccinated animals produced more NK cells on 3 and 10 DPV with minimal effect on B cells (Fig. 2E, F). Overall, IM vaccination appeared to elicit a rapid CD4^+^ T cell and NK cell response, while IN-vaccination resulted in a rapid CD8^+^ T cell response in the lungs.

**Figure 2.**
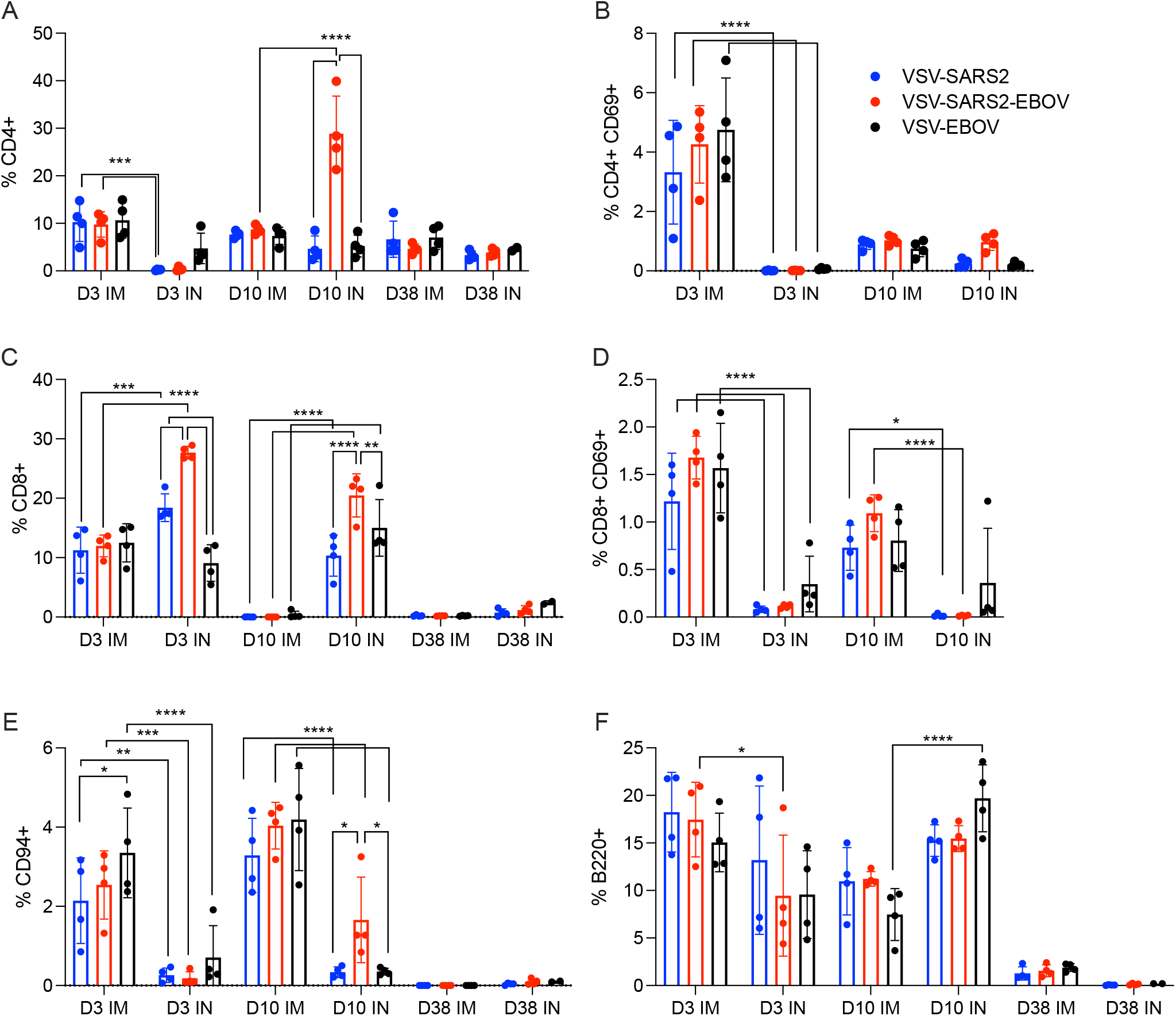
Immunogenicity cellular immune response in the lungs. Single cell lung suspensions were stained for FACS analysis. **(A, B)** CD4^+^ T cells and **(C, D)** CD8^+^ T cells were identified and stained for expression of early activation marker CD69. **(E)** NK cells were identified and stained for expression of CD94. **(F)** B cells were identified and stained for expression of B220. Mean and 95% confidence interval are depicted. Statistical significance determined by two-way ANOVA with Tukey’s multiple comparison is indicated as p<0.0001 (****), p<0.001 (***), p<0.01 (**), and p<0.05 (*).

We examined the same cellular populations in the spleen and in PBMCs of the vaccinated animals. Peak levels of CD4^+^ T cells were measured 10 DPV in the spleen after vaccination by both routes, however, IN vaccination induced more CD4^+^ T cells 38 DPV (Fig. 3A). In contrast, IM vaccination induced more CD8^+^ T cells on 3 and 10 DPV (Fig. 3C). No to limited activated CD4^+^ or CD8^+^ T cell responses were detected (Fig. 3B, D). While IN vaccination resulted in greater numbers of NK cells on 3 and 10 DPV and in more B cells 3 DPV, IM vaccination induced higher numbers of NK cells on 38 DPV and B cells 10 and 38 DPV (Fig. 3E, F). PBMCs of IN-vaccinated animals demonstrated higher levels of CD4^+^ T cells on 38 DPV, while IM vaccination induced significantly more activated CD8^+^ T cells on 10 DPV and CD4^+^ T cells and NK cells on 38 DPV (Fig. S2A-F).

**Figure 3.**
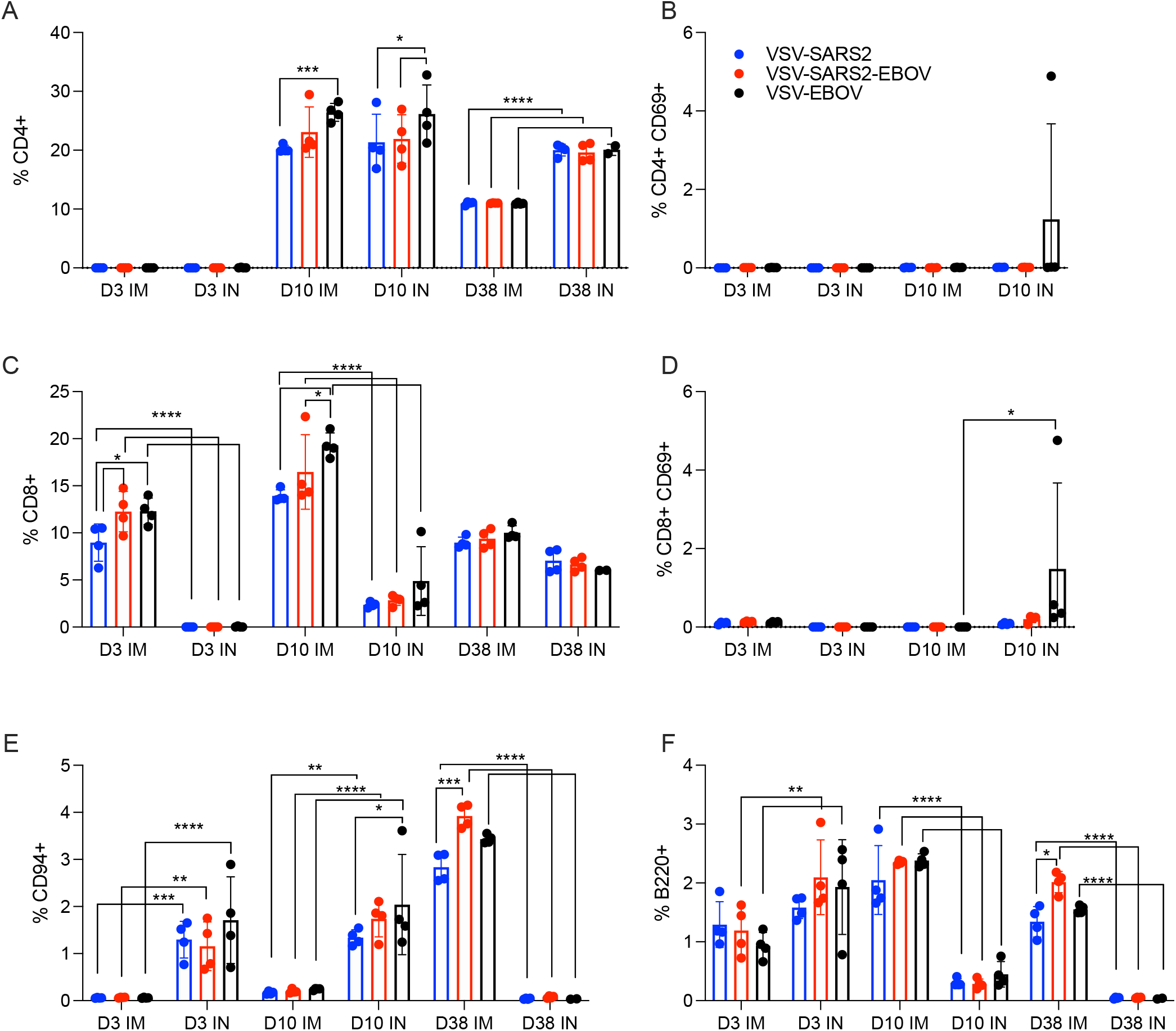
Immunogenicity cellular immune response in the spleen. Single cell splenocyte suspensions were stained for FACS analysis. **(A, B)** CD4^+^ T cells and **(C, D)** CD8^+^ T cells were identified and stained for expression of early activation marker CD69. **(E)** NK cells were identified and stained for expression of CD94. **(F)** B cells were identified and stained for expression of B220. Mean and 95% confidence interval are depicted. Statistical significance determined by two-way ANOVA with Tukey’s multiple comparison is indicated as p<0.0001 (****), p<0.001 (***), p<0.01 (**), and p<0.05 (*).

### VSV-based vaccines protect hamsters from COVID-19 within 10 days

For initial efficacy study in hamsters, we vaccinated groups of 8 animals (4 female and 4 male) with 1×10^5^ PFU either IM or IN with VSV-EBOV (control), VSV-SARS2, or VSV-SARS2-EBOV. The animals were challenged with 1×10^5^ median tissue culture infectious dose (TCID^_50_^) of the SARS-CoV-2 WA1 isolate 28 DPV (day 0) and euthanized 4 days post challenge (DPC) for sample collection. Oral swab samples at the time of necropsy revealed no significant differences in viral shedding as determined by RT-qPCR (Fig. 4A). In contrast, lungs from all vaccinated hamsters presented without lesions (Fig. S3A, B, D, E) and a significant decrease in lung virus loads determined by RT-qPCR (Fig. 4B) and titration (Fig. 4C). All control animals presented with gross lung lesions (Fig. S3C, F) and high lung virus loads (Fig. 4B, C). When we investigated the antibody response 4 DPC, we found higher S-specific IgG titers after both routes of vaccination, however, only titers after IN vaccination were statistically significant (Fig. 4D). Neutralization against the SARS-CoV-2 WA1 isolate revealed significantly higher titers for all vaccinated groups compared to control hamsters (Fig. 4E). In addition, the VSV-SARS2-EBOV vaccine resulted in significantly higher titers after IN administration compared to VSV-SARS2 (Fig. 4E).

**Figure 4.**
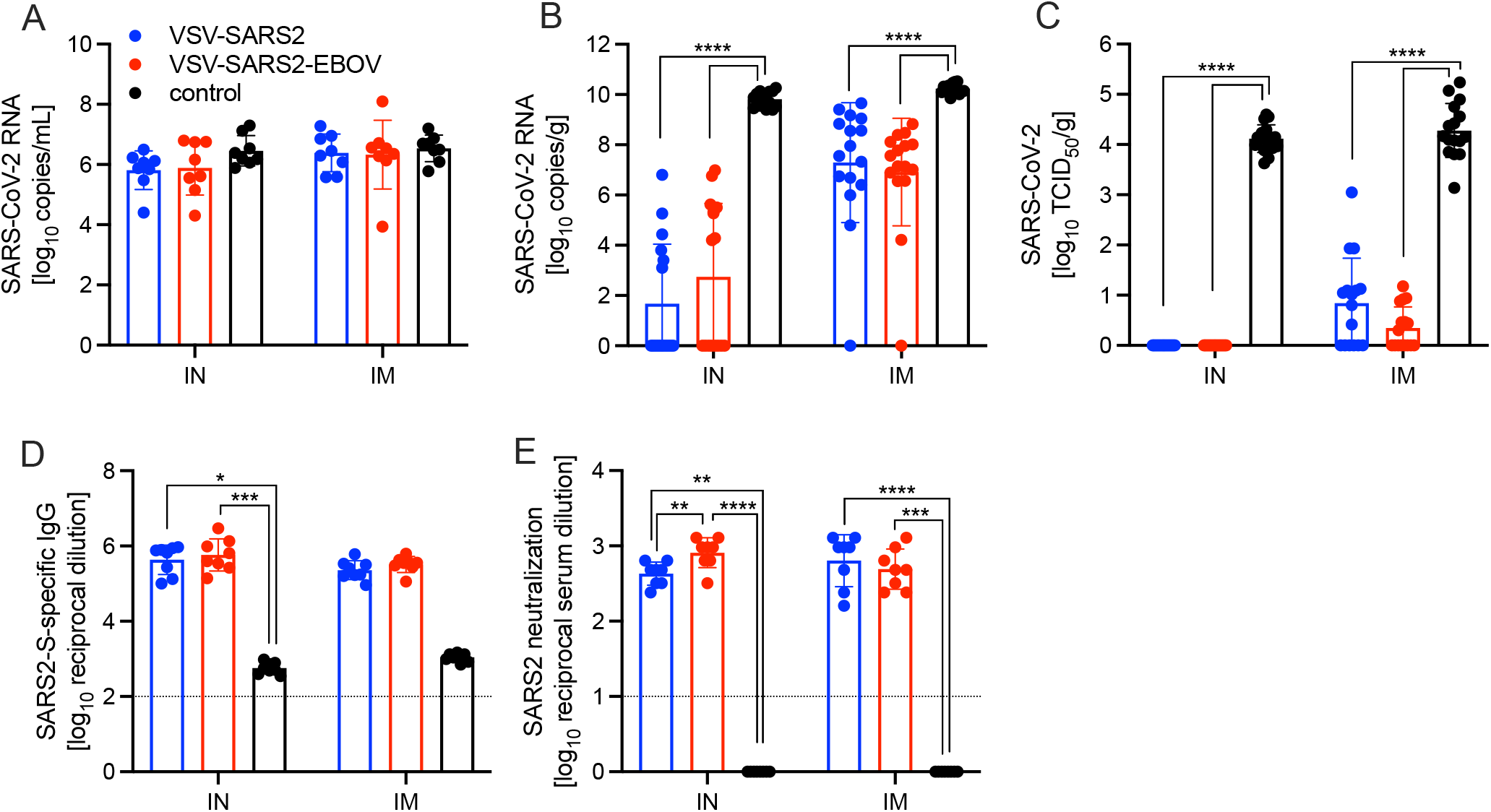
Virus loads and antibody levels in hamsters challenged 28 days post-vaccination. Hamsters were vaccinated with a single dose intramuscularly (IM) or intranasally (IN) 28 days prior to challenge with SARS-CoV-2 WA1. At 4 days after challenge oral swab, lung and serum samples were collected. Levels of SARS-CoV-2 RNA in **(A)** oral swabs and **(B)** lung samples. **(C)** Virus titer in hamster lungs. **(D)** SARS-CoV-2 S-specific IgG and **(E)** neutralizing titers against SARS-CoV-2 WA1 are shown. Geometric mean and geometric SD are depicted. Statistical significance as determined by two-way ANOVA with Tukey’s multiple comparison is indicated as p<0.0001 (****), p<0.001 (***), p<0.01 (**), and p<0.05 (*).

Next, we explored the fast-acting potential of these vaccines by shortening the time between vaccination and challenge to 10 days. Because we did not observe a difference between male and female hamsters in the previous experiment, all following experiments were conducted using female hamsters only. Groups of 6 hamsters were vaccinated with 1×10^5^ PFU with VSV-EBOV (control), VSV-SARS2, or VSV-SARS2-EBOV either IM or IN. The animals were challenged with 1×10^5^ TCID_50_ of the SARS-CoV-2 WA1 isolate 10 DPV (day 0) and euthanized 4 DPC for sample collection. Oral swabs demonstrated a significant decrease in viral RNA indicating reduced shedding in vaccinated animals (Fig. 5A). Gross lung pathology revealed lesions in the control animals (Fig. S3I, L) and, to a lesser extent, in the VSV-SARS2 IM group (Fig. S3G). Hamsters vaccinated with VSV-SARS-EBOV presented without lung lesions (Fig. S3 H, K) as did the VSV-SARS2 IN-vaccinated group (Fig. S3J). Viral loads in the lungs revealed significant differences between vaccinated and control animals by RT-qPCR and virus titration (Fig, 5B, C). Histopathologic analysis of lung samples collected 4 DPC demonstrated evidence of interstitial pneumonia in all control animals (Fig. 6I, K) and was quantified by the application of a pathology score (Fig. 5D). While interstitial pneumonia was significantly reduced in the animals vaccinated IN with both vectors or IM with VSV-SARS2-EBOV (Fig. 5D, 6A, C, G), lung sections of animals in the VSV-SARS2 IM group showed evidence of broncho-interstitial pneumonia consistent with coronaviral pulmonary disease (Fig. 6E). Immunohistochemistry (IHC) revealed the presence of SARS-CoV-2 N in the lungs of control animals only (Fig. 6J, L) indicating control of virus replication in all vaccine groups (Fig. 6B, D, F, H). Analysis of S-specific IgG in the serum of hamsters 4 DPC demonstrated significantly higher S-specific IgG titers after both routes of vaccination (Fig. 5E). Neutralization against the SARS-CoV-2 WA1 isolate revealed significantly higher titers for VSV-SARS2 IN, VSV-SARS2-EBOV IN and VSV-SARS2-EBOV IM vaccine groups compared to the control group (Fig. 5F). In addition, the VSV-SARS2-EBOV vaccine resulted in significantly higher titers after IM administration compared to VSV-SARS2 (Fig. 5E).

**Figure 5.**
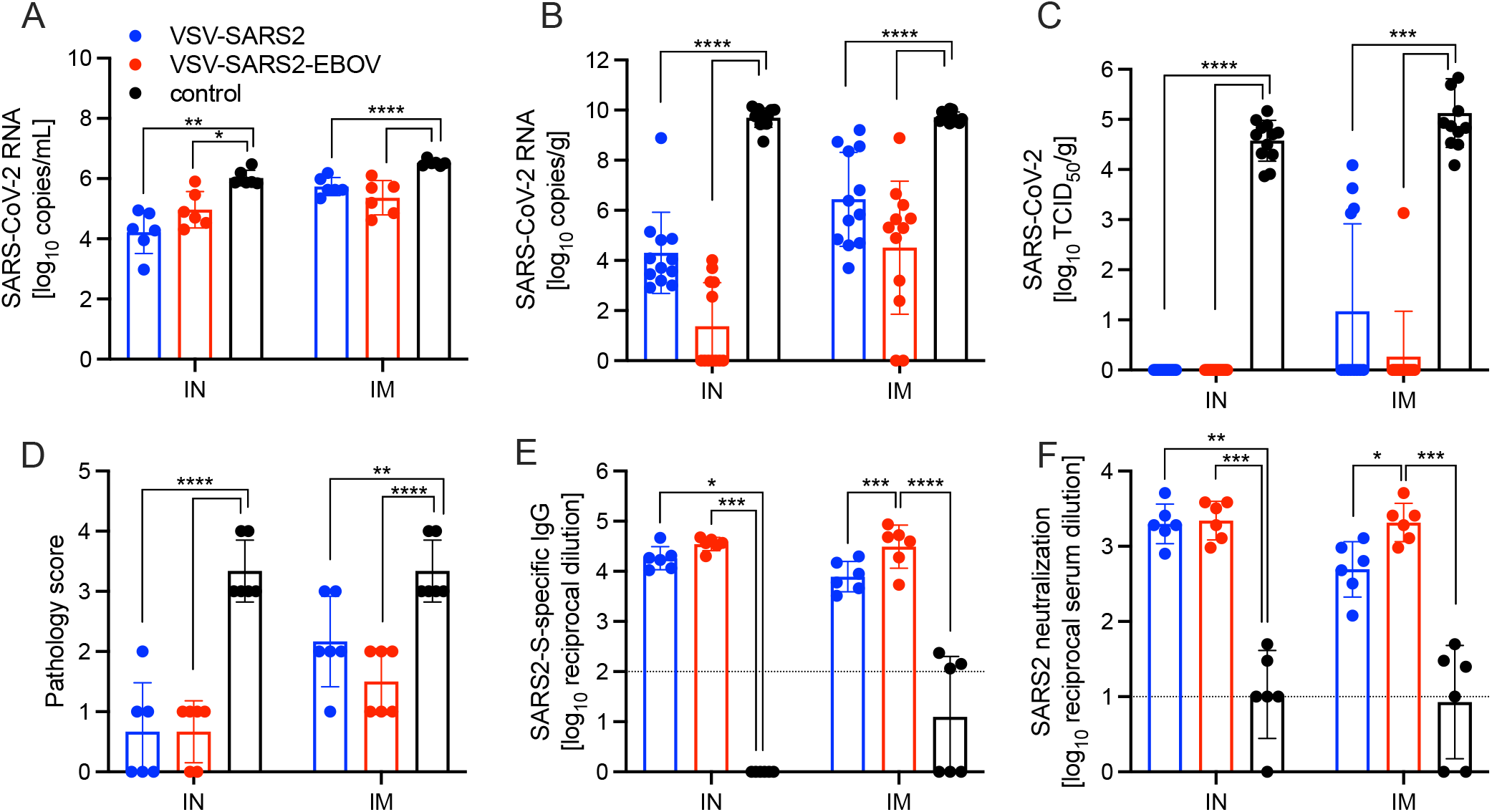
Virus loads and antibody levels in hamsters challenged 10 days post-vaccination (DPV). Hamsters were vaccinated with a single dose intramuscularly (IM) or intranasally (IN) 10 days prior to challenge with SARS-CoV-2 WA1. At 4 days after challenge oral swab, lung and serum samples were collected. Levels of SARS-CoV-2 RNA in **(A)** oral swabs and **(B)** lung samples. **(C)** Virus titer in hamster lungs. **(D)** Lung sections were scored for evidence of interstitial pneumonia. **(E)** SARS-CoV-2 S-specific IgG and **(F)** neutralizing titers against SARS-CoV-2 WA1 are shown. (A-C, E, F) Geometric mean and geometric SD or (D) mean and SD are depicted. Statistical significance as determined by two-way ANOVA with Tukey’s multiple comparison is indicated as p<0.0001 (****), p<0.001 (***), p<0.01 (**), and p<0.05 (*).

**Figure 6.**
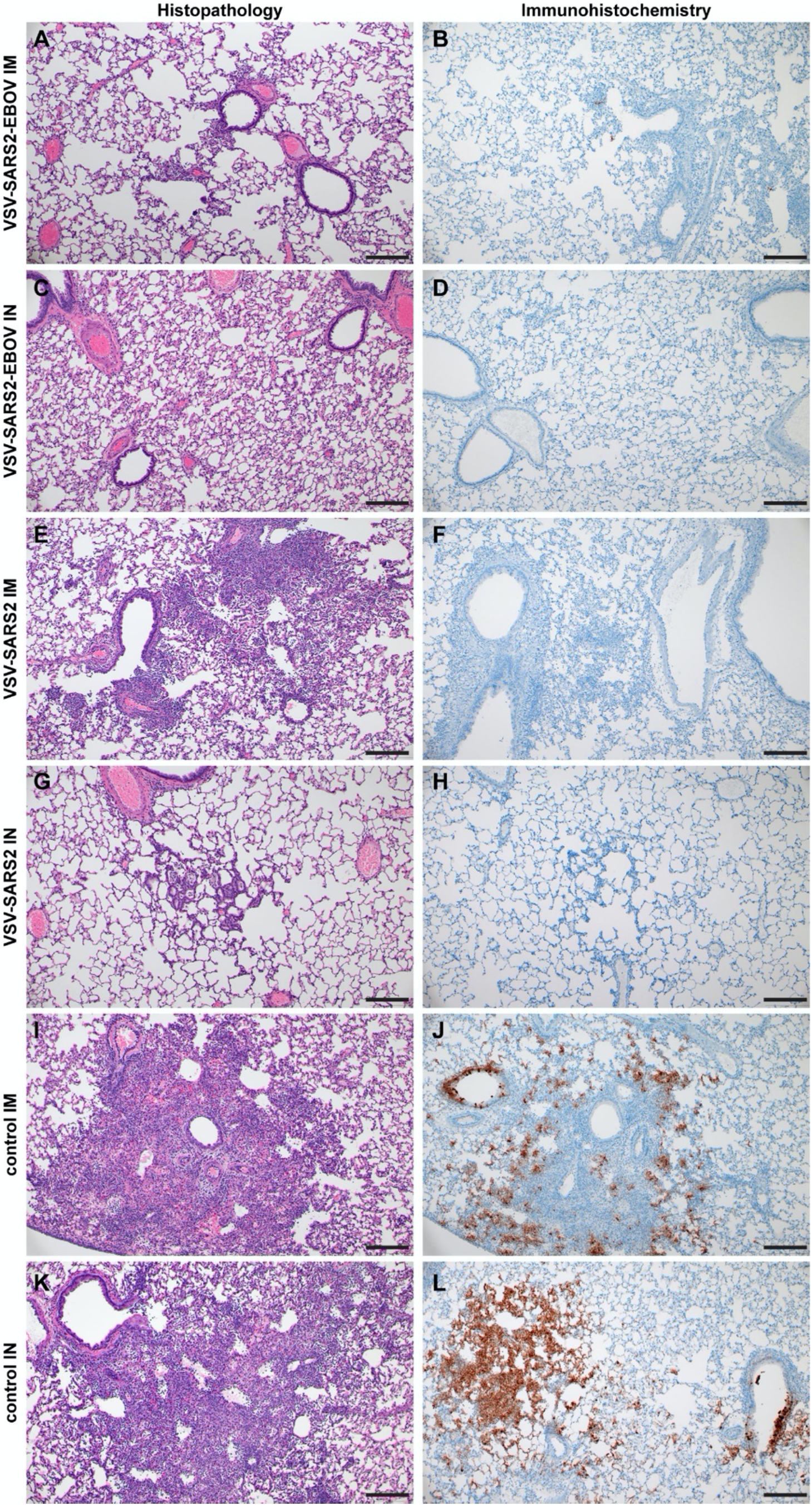
Histopathology and Immunohistochemistry of hamster lungs with challenge 10 DPV. Hamsters were vaccinated 10 days prior to challenge with SARS-CoV-2 WA1. At 4 days after challenge lung samples were collected and stained with H&E or anti-SARS-CoV-2 nucleocapsid (N) antibody for IHC. **(A)** Rare foci of minimal to mild interstitial pneumonia with mild alveolar spillover. **(B)** Rare type I pneumocyte immunoreactivity. **(C)** Lack of notable pulmonary histopathology. **(D)** No immunoreactivity to SARS-CoV-2 N. **(E)** Focus of mild to moderate broncho-interstitial pneumonia with perivascular leukocyte cuffing. **(F)** Limited type I pneumocyte immunoreactivity. **(G)** Rare foci of minimal to mild interstitial pneumonia with type II pneumocyte hyperplasia. **(H)** No immunoreactivity to SARS-CoV-2 N. **(I)** Focus of moderate to severe bronchointerstitial pneumonia with disruption of pulmonary architecture by degenerate and non-degenerate neutrophils, macrophages and cellular debris accompanied with perivascular and pulmonary edema. **(J)** Abundant immunoreactivity to SARS-CoV-2 N in columnar epithelium of bronchioles, type I pneumocytes and alveolar macrophages. **(K)** Moderate broncho-interstitial pneumonia with influx of moderate to numerous leukocytes and limited pulmonary edema. **(L)** Abundant immunoreactivity to SARS-CoV-2 N in bronchiolar epithelium, type I and II pneumocytes and within cellular debris. (200x, bar = 50 μM).

### IN vaccination with VSV-based vaccines protects hamsters against infection with VOC

SARS-CoV-2 VOC are in the focus of efficacy testing for approved vaccines. Therefore, we investigated the protective potential of our VSV-based vaccines against two VOC: B.1.1.7 and B.1.351. Groups of hamsters were vaccinated with 1×10^5^ PFU VSV-EBOV (control), VSV-SARS2, or VSV-SARS2-EBOV (Table S2). VSV-SARS2 IM vaccination was not protective as described above, thus we omitted this group. The hamsters were challenged with 1×10^5^ TCID_50_ of the SARS-CoV-2 B.1.1.7 or B.1.351 10 DPV (day 0) and euthanized 4 DPC for sample collection. Oral swabs taken from vaccinated hamsters showed reduced levels of viral RNA compared to the control groups, however, the differences were not significant for either VOC (Fig. 7A). Gross pathology of the lungs at the time of necropsy 4 DPC revealed lesions in the control groups for both VOC (Fig. S4D, H). Animals in the VSV-SARS2-EBOV IM group presented with limited lung lesions after B.1.1.7 infection (Fig. S4B), whereas VSV-SARS2 or VSV-SARS2-EBOV IN-vaccinated hamsters did not show any lesions grossly (Fig. S4A, C). For the challenge with B.1.351 only hamsters IN-vaccinated with VSV-SARS2 presented with non-lesioned lungs (Fig. S4G). Lung virus loads supported the gross pathology observations for B.1.1.7 challenge with lowest viral RNA detected after IN vaccination (Fig. 7B). Similarly, only IN vaccination significantly reduced SARS-CoV-2 RNA in the lungs of B.1.351-infecetd hamsters albeit to lower extent when compared to B.1.1.7-infecetd hamsters (Fig. 7B). Virus titration of lung samples confirmed the RNA data demonstrating significantly reduced SARS-CoV-2 levels after B.1.1.7 challenge in all vaccinated animals and after B.1.351 challenge in IN-vaccinated hamsters (Fig. 7C). Histopathology revealed significant reductions of evidence of broncho-interstitial pneumonia for IN-vaccinated hamsters with increased vaccination efficacy in the B.1.1.7 group (Fig. 7D). Representative lung sections for each group indicated that VSV-SARS2-EBOV IN vaccination was the most efficacious vaccine against VOC challenge with limited pathological changes and no presence of viral antigen (Fig. S5). Antigen-specific IgG responses were examined from 4 DPC and demonstrated significant titers in all vaccinated hamsters compared to the control groups (Fig. 7E). While there was no significant difference in vaccinated and B.1.1.7-infecetd hamsters for S-specific IgG or neutralization activity (Fig. 7E, F), vaccination with VSV-SARS2-EBOV IN and B.1.351 challenge resulted in a significantly higher S-specific IgG titer (Fig. 7E). Interestingly, this difference could not be confirmed in the neutralization assay. Serum of hamsters vaccinated with VSV-SARS2 IN and challenged with B.1.351 had the highest neutralizing titers against B.1.351 (Fig. 7F). Overall, this data demonstrates that IN vaccination with VSV-based vaccines expressing SARS-CoV-2 S is efficacious against VOC infection within 10 days.

**Figure 7.**
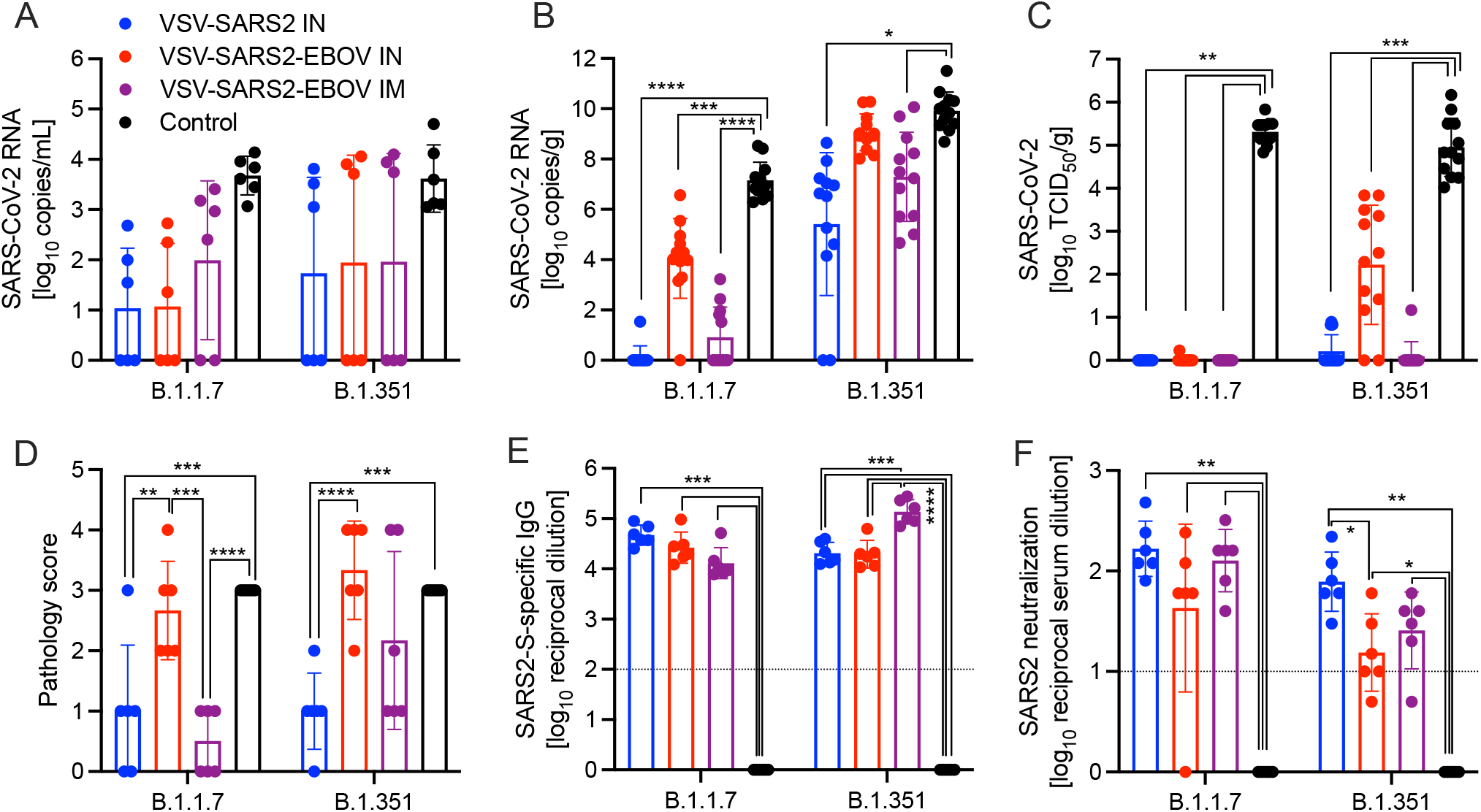
Virus loads and antibody levels in hamsters challenged 10 days post-vaccination with VOC. Hamsters were vaccinated with a single dose intramuscularly (IM) or intranasally (IN) 10 days prior to challenge with SARS-CoV-2 B.1.1.7 or B.1.351. At 4 days after challenge oral swab, lung and serum samples were collected. Levels of SARS-CoV-2 RNA in **(A)** oral swabs and **(B)** lung samples. **(C)** Virus titer in hamster lungs. **(D)** Lung sections were scored for evidence of interstitial pneumonia (1= minimal, 2= mild, 3= moderate, and 4= severe). **(E)** SARS-CoV-2 S-specific IgG and **(F)** neutralizing titers against SARS-CoV-2 WA1 are shown. (A-C, E, F) Geometric mean and geometric SD or (D) mean and SD are depicted. Statistical significance as determined by two-way ANOVA with Tukey’s multiple comparison is indicated as p<0.0001 (****), p<0.001 (***), p<0.01 (**), and p<0.05 (*).

## Discussion

The COVID-19 pandemic is not slowing down and surges in cases caused by VOC are ongoing. The most efficient way to stop the pandemic is vaccination. An effective COVID-19 vaccine would ideally induce protective immunity rapidly after only a single dose, thus reducing the pressure on vaccine production and the healthcare system. Given that VSV-based vaccines often require only a single dose to be effective while inducing a rapid and robust immune response, they offer considerable potential to meet this need. Most of the SARS-CoV-2 vaccines that have been authorized for emergency human use utilize adenovirus- or mRNA-based platforms that require a prime and boost vaccination schedule to fully generate protective immunity (Corbett et al., 2020, Vogel et al., 2020). The prime/boost vaccination strategy requires significant time to achieve full immunity, which intrinsically puts patients at risk. Our goal was to generate a fast-acting single-dose vaccine that could be implemented in an emergency situation for naïve people or as a fast-acting booster vaccination for previously vaccinated individuals or COVID-19 survivors who have waning immunity.

Despite being a live-attenuated vaccine, the VSV vaccine platform has several attributes that contribute to its safety profile, an important consideration for a newly developed vaccine. First, the VSV-based vaccines lack the VSV glycoprotein which is considered the primary virulence factor (Thanh Le et al., 2020). Additionally, VSV is sensitive to interferon α/β and an intact innate immune system is able to control VSV replication (Fathi et al., 2019). Lastly, the VSV-SARS2-EBOV vector is based on the FDA- and EMA-approved EBOV vaccine by Merck and further attenuated by the addition of SARS-CoV-2 S, another safety feature. However, proper toxicity studies for this vector will still be needed for licensure.

The vaccines described here demonstrated strong efficacy regardless of vaccination route when a “classical” 28-day vaccination to challenge model was utilized. When a shorter time to challenge was implemented, the route of vaccination greatly affected the vaccine efficacy with VSV-SARS2 only being effective by the IN route against all three tested viruses. Merck developed a VSV-SARS2 vaccine similar to ours, but recently halted the production because a phase 1 clinical trials demonstrated humoral antigen-specific responses below the levels of COVID-19 survivors following well-tolerated IM administration (Merck & Co., 2021). The report mentions that alternative routes of vaccination including IN are still being investigated, which reflect our data showing increased vaccine efficacy via IN administration. The poor performance of the vaccine may be due to the fact that VSV-SARS2 can only infect ACE2 expressing-cells at the site of vaccination. IN vaccination may be more successful due to the abundant expression of ACE2 in the nasal mucosa compared to fewer in muscle tissue (Sungnak et al., 2020). In contrast, our bivalent VSV-SARS2-EBOV vaccine was effective both after IN and IM administration. This highlights the potential for the use of two glycoproteins with different cellular affinities to promote early replication in different anatomical areas. Interestingly, this is in contrast to the data we have generated in NHPs, where IM VSV-SARS2-EBOV vaccination was more efficacious than IN when 10 days between vaccination and challenge were tested (Furuyama et al., 2021).

The Syrian golden hamster model is a highly susceptible model for SARS-CoV-2, with an ID_50_ of 5 TCID_50_ (Rosenke et al., 2020). Viral RNA and infectious viral titers are high in the respiratory tract of infected hamsters, but do not translate to severe clinical disease manifestations with hamsters displaying minimal weight loss and no to minor outward signs of disease. However, when lung samples are analyzed, histopathology shows evidence of broncho-interstitial pneumonia present in the challenged animals 4 DPC (O’ Donnell et al., 2021, Rosenke et al., 2020). Lung pathology resolved when animals were necropsied 14 and 28 DPC indicating that SARS-CoV-2 infection is a self-limiting disease in this model system (O’ Donnell et al., 2021). The inhibition of severe lung lesions and signs of interstitial pneumonia early during infection is being used as an indicator of vaccine or antiviral therapy efficacy in the hamster model (Rosenke et al., 2020). Histopathologic analysis of lung samples from hamsters vaccinated with either VSV-SARS2 or VSV-SARS2-EBOV demonstrated that regardless of route of immunization, VSV-SARS2-EBOV showed minimal pathological changes. Additionally, no viral antigen was present as shown by IHC. Similarly, lungs of hamsters receiving VSV-SARS2 IN presented with minimal pathological features and no viral antigen was detected. In contrast, lungs of hamsters IM-vaccinated with VSV-SARS2 presented with evidence of interstitial pneumonia and viral antigen was detected within foci of pathology. This led us to conclude that VSV-SARS2-EBOV was a superior vaccine candidate particularly when the vaccine was administered only 10 days prior to challenge.

With the continued emergence of new SARS-CoV-2 VOC harboring mutations that either increase transmissibility or allow for increased evasion from the previously established humoral response, new challenges arise. Existing vaccination strategies and routes of administration must be analyzed to determine the retention of vaccine efficacy against multiple VOC. The two primarily distributed vaccines by Pfizer (BNT162b2) and Moderna (mRNA-1273) have been assessed for sustained efficacy against VOC. A recent report utilizing human serum samples and a pseudovirus neutralization assay determined that vaccination with either mRNA vaccine resulted in moderate decreases in cross-neutralization activity against B.1.1.7. When the cross-reactivity against B.1.351 was assessed the neutralization potential was decreased up to 42.2-(Pfizer) and 27.7-(Moderna) fold, respectively(Garcia-Beltran et al., 2021). In a meta-analysis review both vaccines had various results against B.1.1.7 with a range of 2.6-fold decrease to 3.8-fold increase in live-virus neutralization for the Pfizer vaccine, and a 1.77-fold decrease to 1.6-fold increase for the Moderna vaccine. In contrast, the cross-neutralization of B.1.351 was significantly impacted with 22.8-(Pfizer) and 12.4-(Moderna) fold decreases in live-virus neutralization assays (Noori et al., 2021). The adenovirus-based vaccine from AstraZeneca ChAdOx1 nCOV/AZD1222 followed a similar trend with 70.4-74% percent efficacy against B.1.1.7, but only 10.4-22% efficacy against B.1.351(Knoll and Wonodi, 2021, Harvey et al., 2021, Madhi et al., 2021). Our studies highlight the importance of such experiments. The primary VSV-SARS2-EBOV vector is highly efficacious against the original virus (WA1) within 10 days. When vaccinated hamsters were challenged with the heterologous B.1.1.7 variant the vaccine remained similarly efficacious. However, when the vaccinated hamsters were challenged with the B.1.351 variant, VSV-SARS2-EBOV efficacy was decreased resulting in hamsters’ histopathologic lesions consistent with COVID-19. Interestingly, IN vaccination with VSV-SARS2 was more efficacious against B.1.351 challenge in hamsters within 10 days. However, virus shedding was not reduced after vaccination and challenge similar to other vaccines (Fischer et al., 2021, Wuertz et al., 2021). Future studies will decipher if a longer time between vaccination and challenge results in increased protective immunity after vaccination with both vaccines against challenge with SARS-CoV-2 VOC.

VSV-based vaccines primarily elicit humoral immune response conferring protection from disease (Marzi et al., 2011, Mire et al., 2019, Safronetz et al., 2015). However, we wanted to determine if cellular responses differ based on route of vaccination. The comparison of the immunological characteristics of the different samples assessed in this study is depicted in Fig. S6. The cellular response showed an early CD4^+^ T cell and NK cell response in the lungs of IM-vaccinated hamsters and an early CD8^+^ T cell response in the lungs of IN-vaccinated hamsters. In the spleen, IM vaccination promoted an early CD8^+^ T cell response and a late NK and B cell response, while IN vaccination induced an early NK and B cell response and a late CD4^+^ T cell response. There was little measured activation of T cells in the spleen of vaccinated hamsters and little involvement of neutrophils in either the lung or the spleen. IM vaccination induced a late circulating NK cell response. IN vaccination induced an early circulating CD8^+^ T cell response and a late circulating CD4^+^ T cell and B cell response accompanied by a more robust antibody response. Thus, it appeared that the primary component of the cellular immune response to vaccination with VSV-SARS2 and VSV-SARS2-EBOV is centralized around T cells and NK cells. While the activation marker measured did not show a robust response in the spleen of vaccinated animals, the potential for other effector functions such as stimulation of various cytokines could be present and is an area of interest for future research.

The levels of S-specific IgG measured during the immunogenicity study indicated that both vaccines elicit significantly higher antigen-specific titers compared to control vaccinated animals regardless of vaccination route. This trend also translated to the RBD-specific titers except for 10 DPV in the IM groups. These data indicate that IN vaccination induced a faster and more specific humoral response to potentially neutralizing epitopes. When the functionality of the humoral response was assessed at these time points IN vaccination induced significantly higher neutralization titers 10 and 28 DPV. The humoral response for SARS-CoV-2-S-specific IgG 10 DPV was only significant for VSV-SARS2 when administered IN, while VSV-SARS2-EBOV had significantly higher antigen specific titers compared to control-vaccinated animals regardless of vaccination route. This trend also translated to higher neutralization titers, indicating that not only does VSV-SARS2-EBOV generate higher amounts of antigen-specific antibodies, but also more functional antibodies. The differences in humoral responses were abrogated when hamsters were challenged 28 DPV. The overall humoral response post-challenge compared to vaccination alone elicited a 5-10-fold increase in the response, which may be attributed to the boosting effect of the animals immune system seeing the vaccine antigen for a second time. The overall antigen-specific IgG titers and neutralizing antibody titers 10 and 28 DPV were similar to those reported for ChAdOx1-nCOV/AZD1222 when administered as a single dose 28 days prior to challenge in the hamster model. With the limited immunological tools available for the hamster model the route of vaccination dictates the skew of the cellular response for either vaccine. The overall humoral response is stronger for the VSV-SARS2-EBOV vaccine, which is reflective of the pathologic findings. Traditionally VSV vaccination has been more reliant on a strong humoral response to mediate protection, which leads us to conclude that the differences in SARS-CoV-2-S-specific IgG and neutralization titers are of more importance than the difference in the cellular changes due to the route of vaccination.

Taken together, we generated two effective, single-dose vaccines against COVID-19 efficacious within 10 days in a Syrian golden hamster vaccine-challenge model. VSV-SARS2-EBOV is effective 28 and 10 DPV, regardless of route of vaccination. Our results suggest that IN is the optimal route of vaccination in the hamster model for VSV-based vaccines as well as other vaccines (van Doremalen et al., 2021). Future studies will address the impact of preexisting immunity to SARS-CoV-2 S or EBOV GP in our vaccine, however, we do not anticipate a major effect as both antigens are able to drive the replication of the vaccine virus (Kirby, 2021, CDC, 2021). Furthermore, we will investigate the addition of another SARS-CoV-2 antigen into the vaccine to promote a stronger T cell response, as these responses are typically longer lasting. At this time, the VSV vaccines presented here have a high potential as a boosting option after the already approved vaccines due to their fast-acting potential and the elicitation of primarily a humoral response in contrast to the predominantly T cell-driven immune response after adenovirus- and mRNA-based vaccination (Corbett et al., 2020).

## Acknowledgments

We thank the Rocky Mountain Veterinary Branch, NIAID for supporting the animal studies, and Anita Mora (NIAID) for assistance generating the pathology figures. We also thank members of the Molecular Pathogenesis Unit, Virus Ecology Section, and Research Technology Branch (all NIAID) for their efforts to obtain and characterize the SARS-CoV-2 isolates.

## Author contributions

A.M. conceived the idea and secured funding. K.L.O. and A.M. designed the studies. K.L.O., C.S.C., A.J.G., C.M.L., W.F., and A.M. conducted the studies. K.L.O., K.S., T.G., T.T., W.F. and A.M. processed the samples and acquired the data. K.L.O., C.S.C., and A.M. analyzed and interpreted the data. K.L.O., C.S.C., and A.M. prepared the manuscript. All authors approved the manuscript.

## Supplementary Materials

Table S1. Hamster group sizes used in this study.

Table S2. Flow cytometry antibodies for hamster samples.

Figure S1. Schematic and characterization of VSV-based vaccines.

Figure S2. Vaccine-induced cellular immune response in PBMCs.

Figure S3. Hamster lung gross pathology after vaccination and challenge with SARS-CoV-2 WA1.

Figure S4. Hamster lung gross pathology after vaccination and challenge with SARS-CoV-2 VOC.

Figure S5. Histopathology and Immunohistochemistry of hamster lungs with VOC challenge 10 DPV.

Figure S6. Schematic presentation of the different immune cells responding to IM and IN vaccination using VSV-based vaccines.

