## supplement for "Optimization of single dose VSV-based COVID-19 vaccination in hamsters"

**Supplemental Materials**

Table S1. Hamster group sizes used in this study.
Table S2. Flow cytometry antibodies for hamster samples.
Figure S1. Schematic and characterization of VSV-based vaccines.
Figure S2. Vaccine-induced cellular immune response in PBMCs.
Figure S3. Hamster lung gross pathology after vaccination and challenge with SARS-CoV-2 WA1.
Figure S4. Hamster lung gross pathology after vaccination and challenge with SARS-CoV-2 VOC.
Figure S5. Histopathology and Immunohistochemistry of hamster lungs with VOC challenge 10 DPV.
Figure S6. Schematic presentation of the different immune cells responding to IM and IN vaccination using VSV-based vaccines.

**Table S1. Hamster group sizes used in this study.**

| Group | VSV-SARS2 IN/IM | VSV-SARS2-EBOV IN/IM | Control IN/IM |
|---|---|---|---|
| 3 DPV | 4/4 | 4/4 | 4/4 |
| 10 DPV | 4/4 | 4/4 | 4/4 |
| 21 DPV | 6/6 | 8/8 | 8/8 |
| 38 DPV | 4/4 | 4/4 | 3/3 |

| WA challenge | VSV-SARS2 IN/IM | VSV-SARS2-EBOV IN/IM | Control IN/IM |
|---|---|---|---|
| 28 DPV | 8/8 | 8/8 | 8/8 |
| 10 DPV | 6/6 | 6/6 | 6/6 |

| 10 DPV variants | VSV-SARS2 IN | VSV-SARS2-EBOV IN/IM | Control IN/IM |
|---|---|---|---|
| B.1.1.7 | 6 | 6/6 | 3/3 |
| B.1.351 | 6 | 6/6 | 3/3 |

VSV vesicular stomatitis virus; SARS2 SARS-CoV-2; EBOV Ebola virus; IN intranasal; IM intramuscular; DPV days post-vaccination; WA SARS-CoV-2 Washington isolate (original strain).

**Table S2. Flow cytometry antibodies for hamster samples.**

| Ab target | clone | Ab target | clone | Ab target | clone |
|---|---|---|---|---|---|
| CD3 | 17A2 | Ly6C | HK1.4 | Ki67 | 16A8 |
| CD3 | KT3.1.1 | **Ly6G** | **1A8** | CD19 | 6D5 |
| Cd49b | Dx5 | **CD11b** | **M1/70** | CD27 | LG.3A10 |
| **CD69** | **H1.2F3** | CD11c | HL3 | CD138 | 281-2 |
| CD56 | 809220 | **CD4** | **GK1.5** | IL-2 | JES6-1A12 |
| Granzyme B | QA16A02 | **CD8** | **341** | IL-4 | 11B11 |
| CD107a | 1D4b | CD44 | IM7 | IA/IE | M5/114.15.2 |
| CD45 | 30-F11 | IFN-$\gamma$ | XMG1.2 | **CD94** | **18d3** |
| **Live/Dead** | | IL-17a | TC11-18H10.1 | IA/IE | 14-4-4S |
| **B220** | **RA3-6B2** | CD3 | 145-2C11 | | |

Ab antibody. All antibodies were tested at dilutions ranging from 1:50 to 1:1,600. Antibodies highlighted in red font were successfully used in our assays.

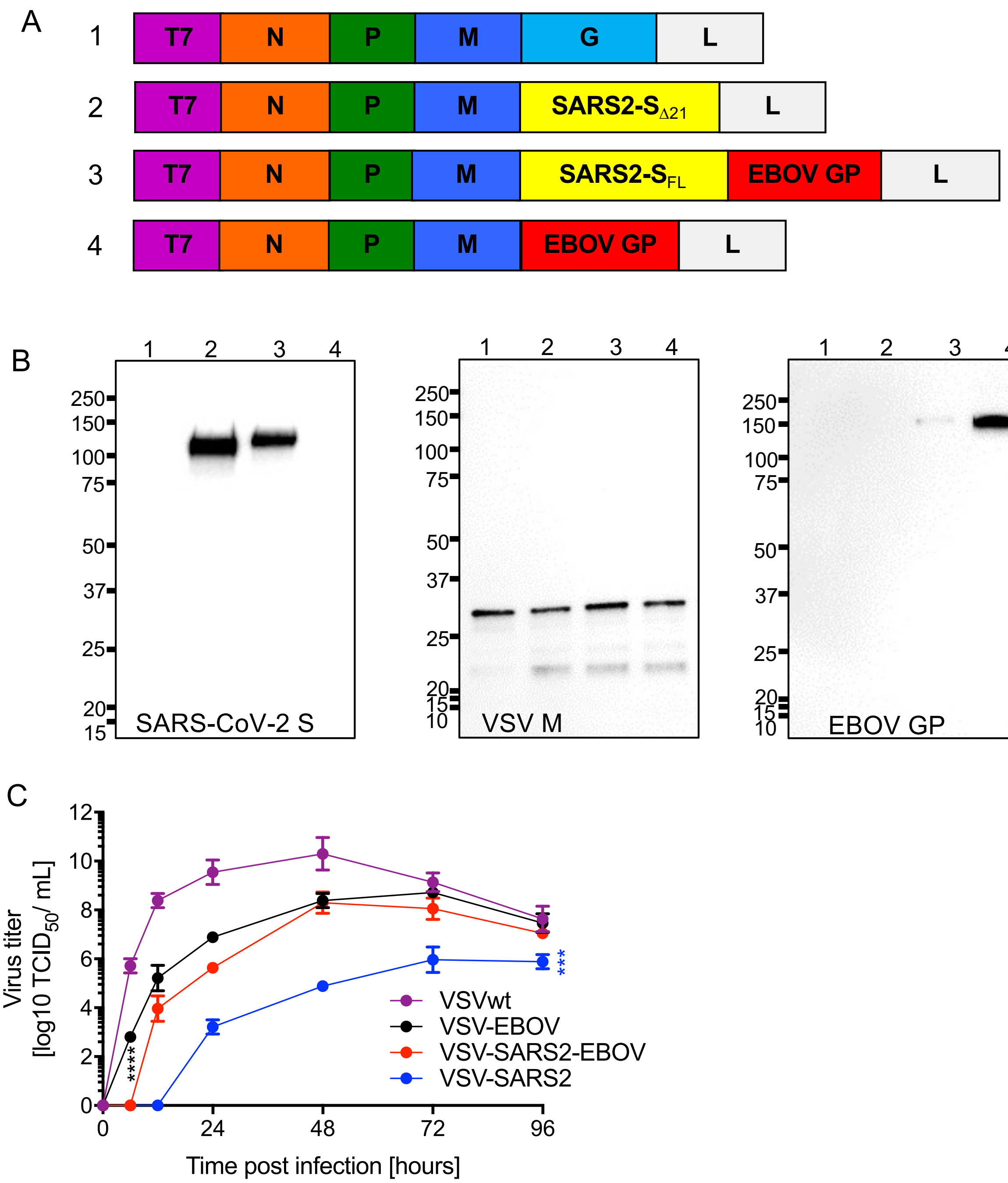


**Figure S1. Schematic and characterization of VSV-based vaccines. (A)** Schematic illustrating vaccine vector design. T7 promotor; N nucleoprotein; P phosphoprotein; M matrix protein; EBOV GP Ebola virus glycoprotein; L RNA-dependent RNA polymerase; SARS2-S SARS-CoV-2 S. **(B)** Western blot analysis of cell supernatant samples containing VSV vaccines probed for SARS-CoV-2 S (left), VSV M (middle) or EBOV GP (right). 1 VSV wildtype (VSVwt); 2 VSV-SARS2; 3 VSV- SARS2-EBOV; 4 VSV-EBOV. **(C)** Viral growth kinetics on Vero E6 cells. Geometric mean and geometric SD are depicted. Statistical significance as determined by two-way ANOVA with Tukey's multiple comparison is indicated as $p<0.0001$ (****), and $p<0.001$ (***).

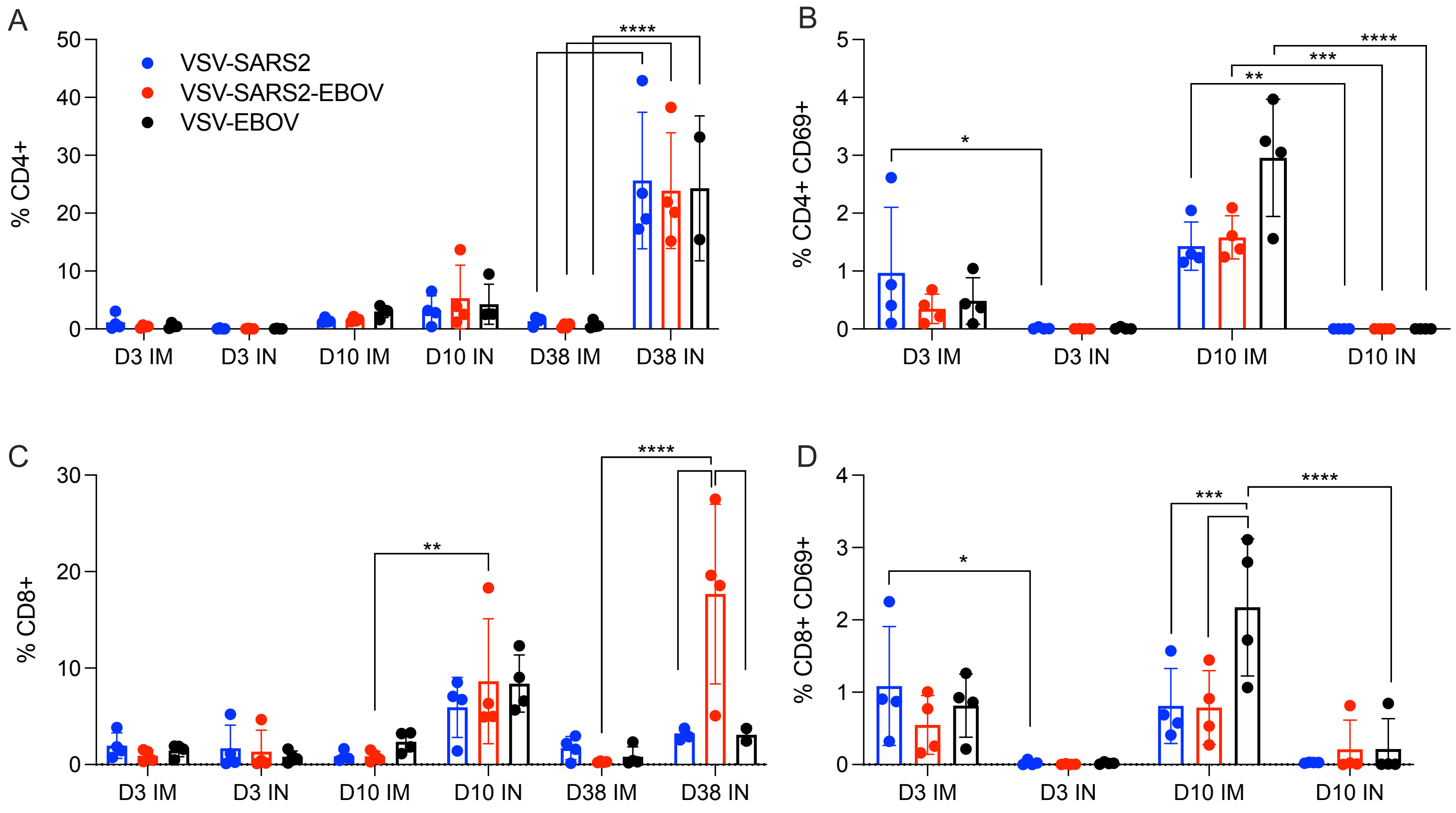


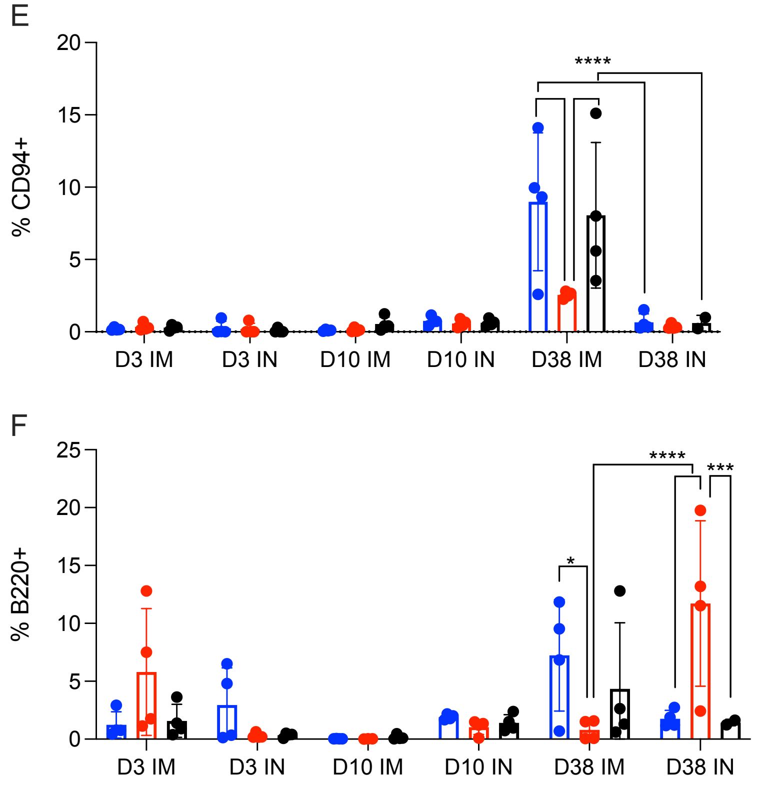


**Figure S2. Vaccine-induced cellular immune response in PBMCs**. Single cell PBMCs suspensions were stained for FACS analysis. **(A, B)** $CD4^+$ T cells and **(C, D)** $CD8^+$ T cells were identified and stained for expression of early activation marker CD69. **(E)** NK cells were identified and stained for expression of CD94. **(F)** B cells were identified and stained for expression of B220. Mean and SD are depicted. Statistical significance as determined by two-way ANOVA with Tukey's multiple comparison is indicated as p<0.0001 (****), p<0.001 (***), p<0.01 (**), and p<0.05 (*).

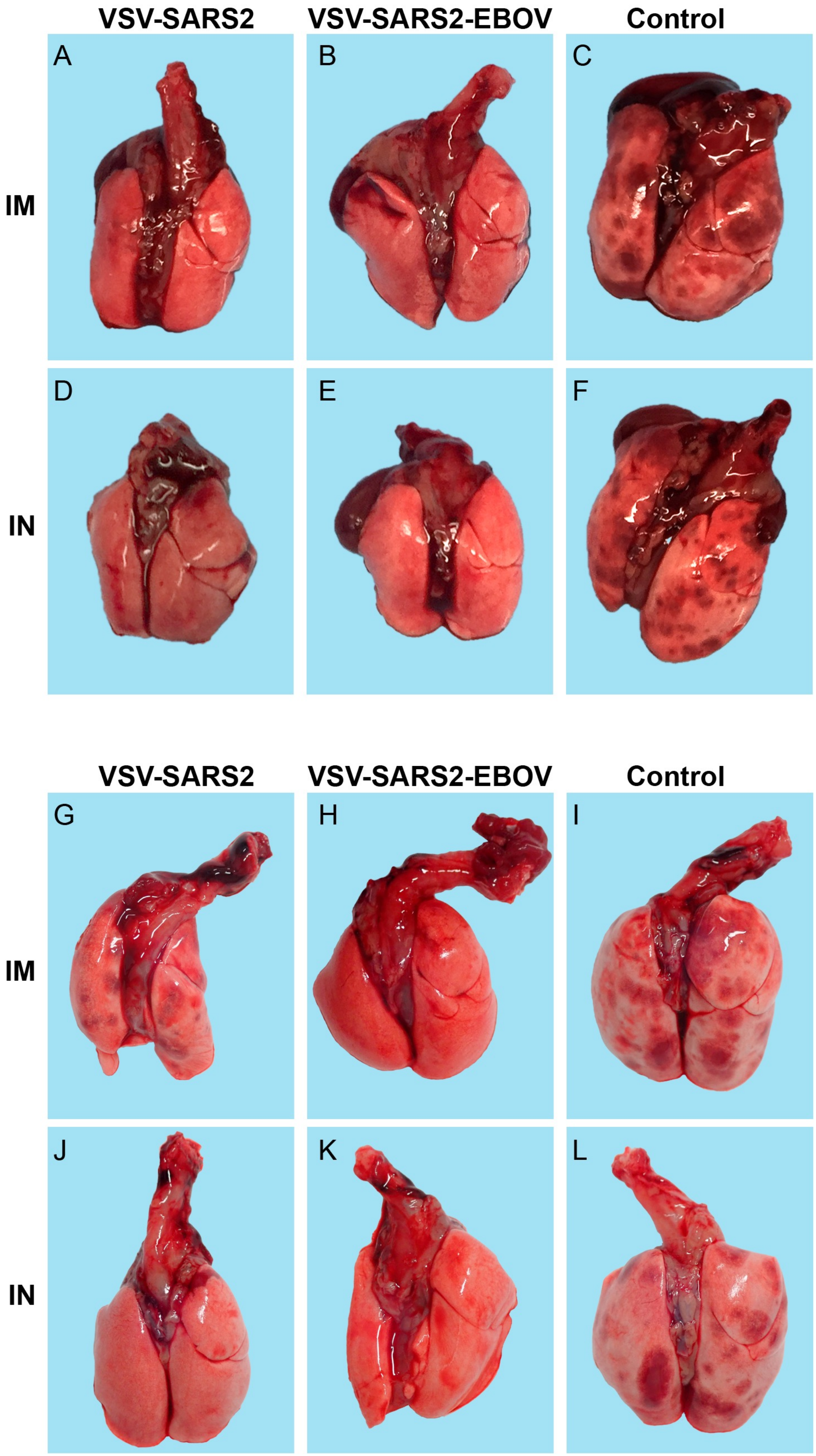


**Figure S3. Hamster lung gross pathology after vaccination and challenge with SARS-CoV-2 WA1.** Groups of hamsters were vaccinated with a single dose of the indicated vaccine by either the intramuscular (IM) or intranasal (IN) route. Challenge occurred (A-F) 28 days or (G-L) 10 days after vaccination. Representative pictures of hamster lungs with lesions for each vaccine group at 4 days post-challenge.

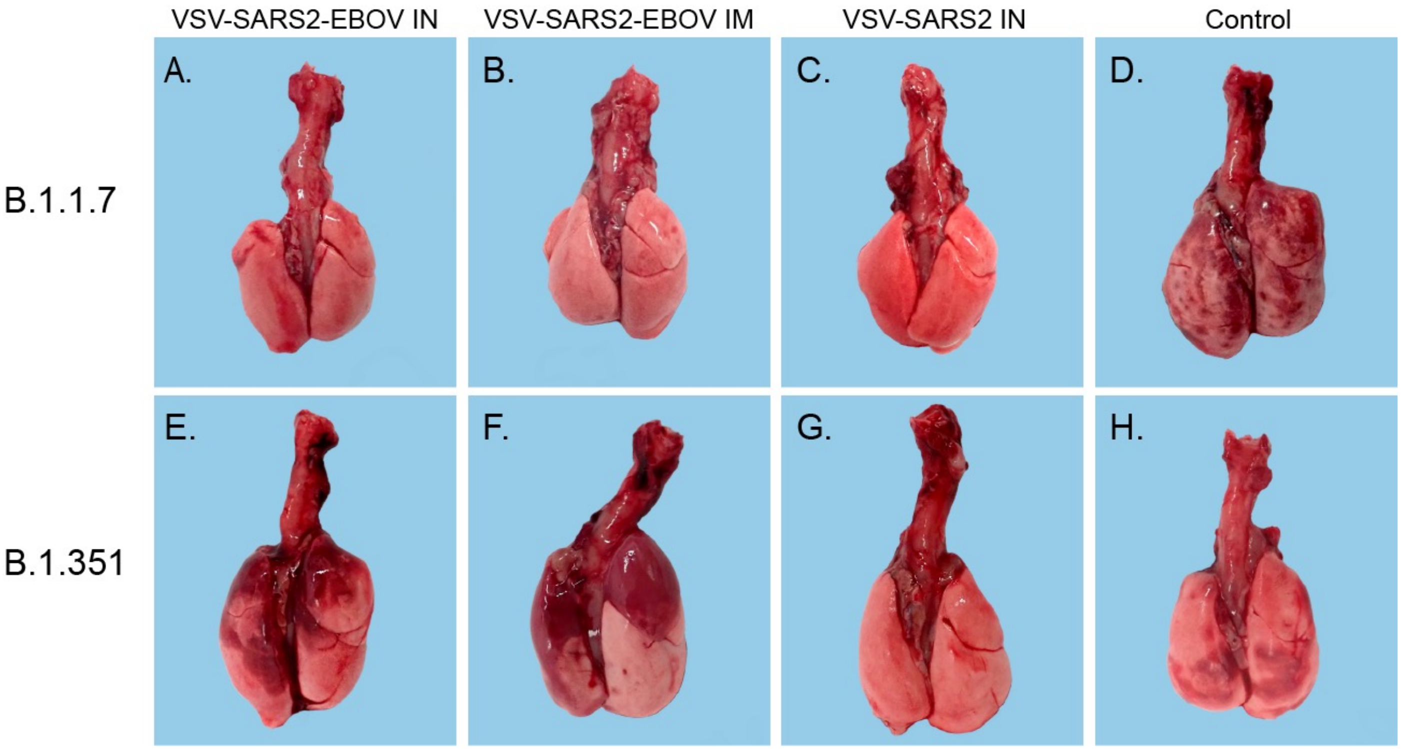


**Figure S4. Hamster lung gross pathology after vaccination and challenge with SARS-CoV-2 VOC.** Groups of hamsters were vaccinated with a single dose of the indicated vaccine by either the intramuscular (IM) or intranasal (IN) route. Challenge occurred 10 days after vaccination with SARS-CoV-2 B.1.1.7 or B.1.351. Representative pictures of hamster lungs with lesions for each vaccine group at 4 days post-challenge.

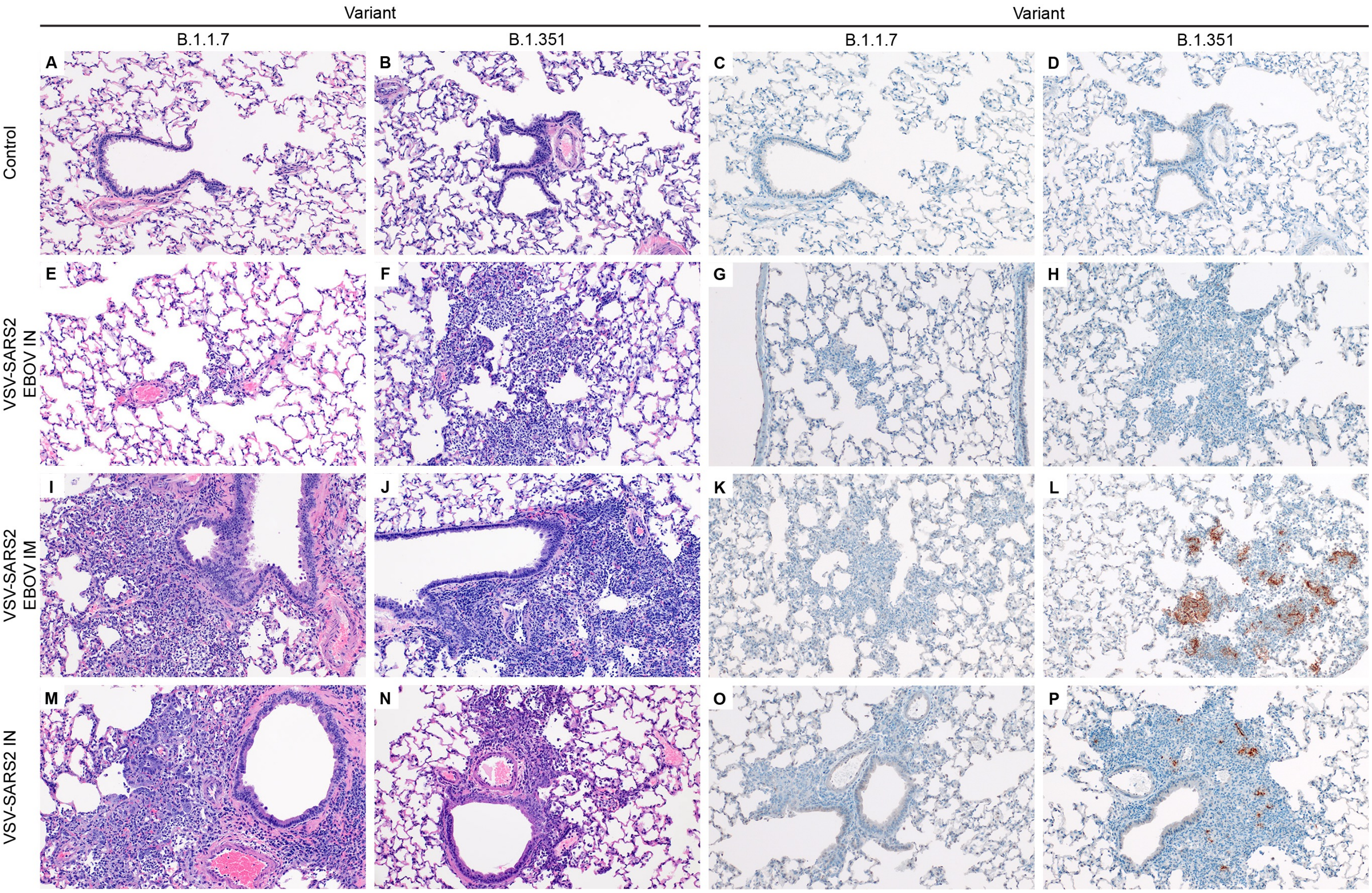


**Figure S5. Histopathology and Immunohistochemistry of hamster lungs with VOC challenge 10 DPV.** Hamsters were vaccinated 10 days prior to challenge with SARS-CoV-2 B.1.1.7 or B.1.351. At 4 days after challenge lung samples were collected and stained with H&E (A-B,E-F,I-J,M-N) or anti-SARS-CoV-2 nucleocapsid (N) antibody for IHC (C-D,G-H,K-L,O-P). All images are 200x. **(A, B)** Normal pulmonary architecture in a mock-infected hamster. **(C, D)** Normal pulmonary architecture showing lack of SARS-CoV-2 immunoreactivity. **(E)** Rare foci of interstitial pneumonia are observed in VSV-SARS2-EBOV intranasal (IN)-vaccinated hamsters when challenged with the B.1.1.7 variant. **(F)** Rare foci of minimal to mild interstitial pneumonia are observed in VSV-SARS2-EBOV IN-vaccinated hamsters when challenged with the B.1.351 variant. **(G, H)** Lack of SARS-CoV-2 immunoreactivity in foci of minimal to mild interstitial pneumonia in the VSV-SARS2-EBOV IN-vaccinated hamsters following challenge with either B.1.1.7 or B.1.351. **(I, J)** Mild to moderate interstitial pneumonia focused on terminal airways are observed in VSV-SARS2-EBOV intramuscular (IM)-vaccinated hamsters when challenged with B.1.1.7 and B.1.351 variants. **(K)** Lack of SARS-CoV-2 immunoreactivity in foci of interstitial pneumonia in a VSV-SARS2-EBOV IM-vaccinated hamster when challenged with B.1.1.7. **(L)** Abundant SARS-CoV-2 immunoreactivity is observed in type I and II pneumocytes with fewer macrophages in a VSV-SARS2-EBOV IM-vaccinated hamster challenged with B.1.351. **(M, N)** Mild to moderate interstitial pneumonia are observed adjacent to bronchioles and terminal airways in the VSV-SARS2 IN vaccinated hamsters when challenged with either the B.1.1.7 or B.1.351 variants. **(O)** Lack of SARS-CoV-2 immunoreactivity in foci of interstitial pneumonia in a VSV-SARS2 IN-vaccinated hamster when challenged with B.1.1.7. **(P)** Rare SARS-CoV-2 immunoreactivity is observed primarily in type I and II pneumocytes in a VSV-SARS2 IN-vaccinated hamster after B.1.351 challenge.

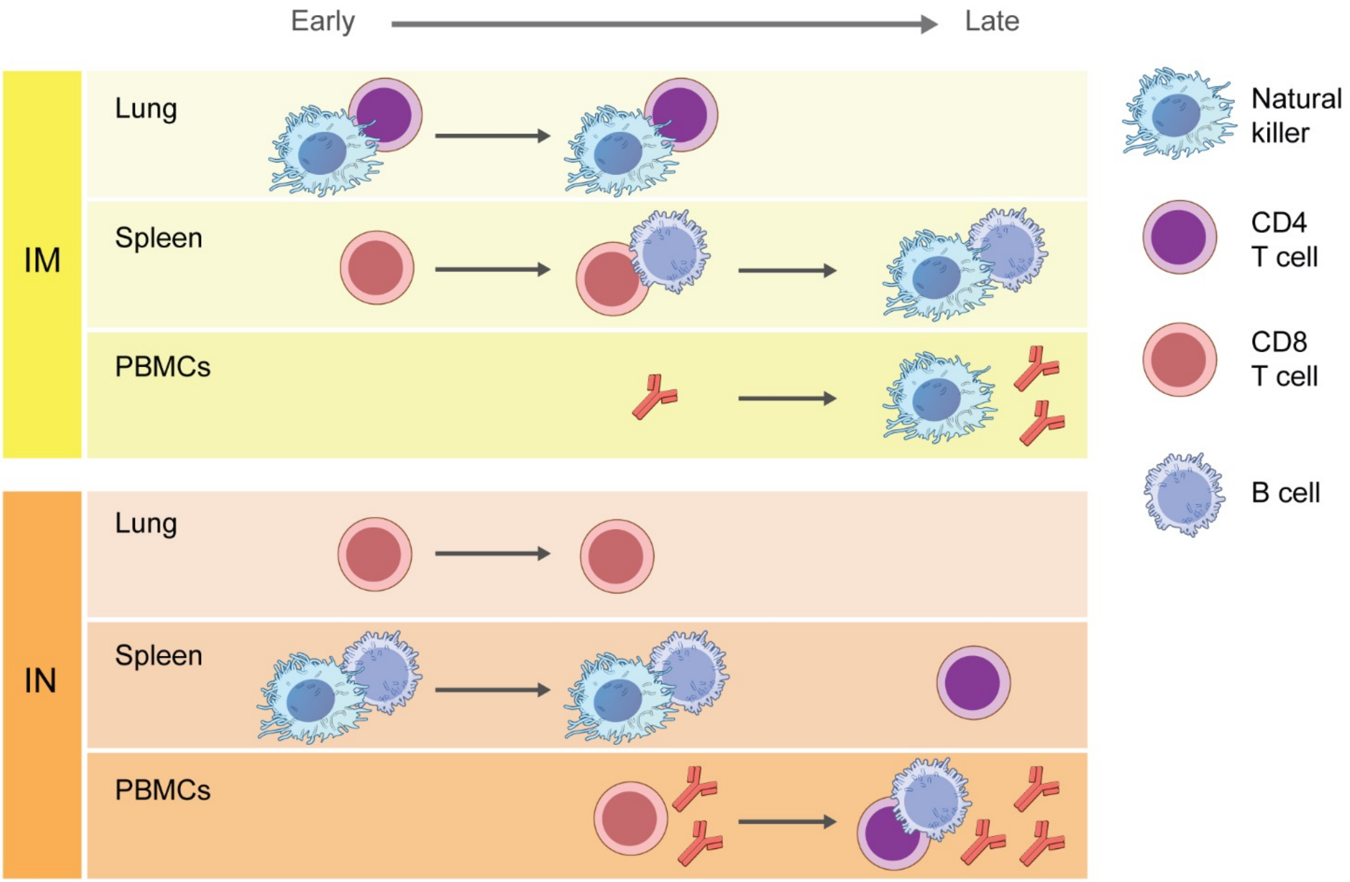


**Figure S6. Schematic presentation of the different immune responses to IM and IN vaccination using VSV-based vaccines.** This diagram depicts the differences in cellular and humoral immune responses in three different anatomical locations. The picture also shows the change/maturation over time highlighting a clear shift in immunological phenotypes. IM intramuscular; IN intranasal; PBMCs peripheral blood mononuclear cells.
